# Postprandial hypertriglyceridemia activates microglia to promote satiety via an ASC dependent signaling

**DOI:** 10.64898/2026.10.04.756524

**Authors:** Juliette Salvi, Emmanuelle Nédélec, Lucie Leret, Fabienne Liénard, Tatiana Landré, Selma Ben Fradj, Valentin Garcia, Vincent Gigot, Bruno Patris, Vivien Pires, Arnaud Polizzi, Clara Sanchez, Thomas Gautier, Hervé Guillou, Tony Heurtaux, Virginie Petrilli, Clémence Blouet, Michael Heneka, François Rassendren, Carole Rovère, Etienne Audinat, Hélène Hirbec, Vincent Compan, Doriane Trompier, Alexandre Benani

## Abstract

Reactive microglia are commonly considered neurotoxic and detrimental for brain functioning. Chronic consumption of a high-fat diet strongly activates microglia, disrupts the central control of energy metabolism, contributes to obesity, and promotes cognitive decline. In this study we report an unexpected beneficial function of microglia reaction at the time scale of a meal, whereby they promote satiety in a macronutrient-dependent manner. We show that murine microglia react to ingested dietary fat within three hours. Specifically, postprandial hypertriglyceridemia induced microglial hyper-ramification alongside with a transcriptional reprogramming promoting communication with neurons. This effect occurs in discrete brain areas, including the hypothalamus and the brainstem, which control hunger. Remarkably, microglia-specific genetic ablation of Apoptosis-associated Speck-like protein containing a CARD (named ASC), a master regulator of metabolism, immune function and cell shape, profoundly alters satiety following high-fat food consumption. These results reveal that microglia engage an immune signaling pathway to rapidly alert the neural circuits and activate an anorectic response that limits overconsumption of dietary fat. Unexpectedly, this regulatory process is sexually dimorphic, present in males only. Furthermore, we found that the postprandial microglial reaction is bypassed during obesity, providing a new therapeutic target to treat maladaptive eating disorders.

## Main

In everyday life, individuals require a continuous supply of energy to sustain physical and mental activities, and to maintain the integrity of the body. Daily food intake properly compensates energy expenditures. As a result, adiposity, which serves as the main energy store, and body weight remain relatively stable during adulthood ^1^. This continuous regulation over time is referred to as energy homeostasis. The brain is at the core of this homeostatic process, operating within a neuroendocrine loop ^2,3^. Discrete neuronal populations are specialized in sensing of energy availability ^4^. One of the best characterized homeostatic neurocircuit controlling food intake is the melanocortin system ^5^. It includes POMC neurons, which express the pro-opiomelanocortin (POMC) peptide and promote satiety during positive energy balance, and NPY/AgRP neurons, which co-express neuropeptide Y (NPY) and agouti-related protein (AgRP), and stimulate food seeking when energy stores and nutrients availability are low ^6,7^. These two antagonistic neuronal populations are located in the arcuate nucleus of the hypothalamus, a small ventral brain region, with permeable vasculature that allows circulating molecules such as nutrients and hormones to reach the parenchyma ^8^.

Microglia, the resident immune cells of the brain, are critical regulators of the organization and activity of the melanocortin system ^9–12^. Usually, these cells exhibit fine and ramified branches oriented radially from a small soma. When mice are kept on high-fat diet (HFD), microglia undergo drastic morphological and functional changes, increasing their volume ^11,13–15^, and adopting an inflammatory phenotype ^13,14,16–18^. Inhibition of the inflammatory signaling in microglia reduces HFD-induced obesity ^19^. Inversely, chronic activation of pro-inflammatory signaling in microglia causes hyperphagia and induces body weight gain. During chronic HFD, reactive microglia also promote compulsive eating behavior by modulating neurons located in the anterior paraventricular thalamus, a reward center ^20^. Thus, HFD-induced microglial reactivity is currently viewed as a pathogenic process that worsens metabolic diseases ^21,22^. However, reactive microglia do not only exhibit detrimental effects on energy homeostasis. Lipoprotein lipase (LPL), a key enzyme mediating the cellular uptake of lipids from circulating lipoproteins, is highly expressed in microglia. Mice with LPL-deficiency in microglia display impaired microglial reactivity and increased vulnerability to diet-induced obesity. These findings demonstrated that microglial metabolism is crucial for maintaining energy homeostasis upon caloric pressure ^23^. Moreover, activating microglia using chemogenetics improves blood glucose control during chronic consumption of HFD ^24^.

Whether microglia are involved in the regulation of energy balance in healthy condition is still a matter of debate ^25^. Yet, microglia can directly sense nutrients and metabolic hormones, a key step in the brain control of energy homeostasis ^26^. Sensory stimuli that derived from foods in the oral cavity are also integrated by microglia within minutes ^27^, thereby contributing to the cephalic phase that amplifies postprandial insulin release ^27^. Moreover, hypothalamic microglia change their cell volume within few hours after HFD ingestion ^28^, suggesting that microglia are involved in the short-term control of energy balance.

In the current study in mice, we demonstrate that brain microglia undergo rapid morphological transitions after a meal in tandem with a transcriptomic reprogramming indicative of enhanced microglia-neuron communication. This postprandial microglial plasticity is characterized by a marked hyper-ramification. The response is fast, occurring within 3 hours after eating, and reversible, resolving within 24 hours. It is macronutrient-dependent, being triggered by high-fat but not standard chow. It is also region-specific, occurring in few discrete brain areas that control appetite, i.e. the *arcuate nucleus* in the hypothalamus, and the *nucleus tractus solitarii* in the brainstem, and not throughout the entire brain. Notably, this response is sexually dimorphic, developing in males and not in females. Selective microglial deletion of the Asc gene, a regulator of cytoskeletal dynamics and immune functions, totally abolishes this postprandial microglial plasticity and leads to overfeeding, revealing the adaptive value of the response. Interestingly, this physiological mechanism is lost in obese hyperphagic mice. Together, these findings identify microglia as critical regulators of food intake during normal physiology.

## Results

### Consumption of HFD for 3 hours elicits hyper-ramification of microglia in the arcuate nucleus of the hypothalamus

To explore whether microglial structure varies at the time scale of a meal, we investigated microglia morphology in the brain of 8/9-week-old male C57BL/6J mice under different prandial states. For that, mice were assigned to three experimental groups, including one preprandial group and two postprandial groups that were fed either with SD or HFD for 3 hours, respectively. To synchronize feeding behavior and retain normal physiology, mice were subjected to 2-hour food restriction prior to the dark period, and food was given *ad libitum* at light off, at the beginning of the active phase (Fig. 1a). With this paradigm, mice ate approximatively 0.9 and 1.2 grams of SD and HFD respectively during the 3h-period (Fig. 1b). In addition, HFD consumption but not SD caused postprandial hypertriglyceridemia (Fig. 1c). We performed post-mortem histological examination of Iba1 immunostained myeloid cells in fixed brain slices from preprandial, 3h-SD and 3h-HFD mice. In contrast to the microglial specific marker P2RY12 ^29,30^, Iba1 marker enables accurate detection of microglia shape, including cell body and processes (Extended Data Fig. 1a). Among all detected Iba1 cells in the hypothalamus, 100% were also labeled with P2RY12, confirming that hypothalamic Iba1 positive cells were *bona fide* microglia (Extended Data Fig. 1a). Quantitative morphometric three-dimensional analysis revealed longer processes and increased branching of Iba1 positive microglia located in the arcuate nucleus of the hypothalamus (ARC) after 3h-HFD compared to preprandial and 3h-SD conditions (Fig. 1d,e). No difference in microglial morphology was detected between 3h-SD and preprandial mice. Similar morphological changes occurred in the brainstem, in particular in the nucleus of the solitary tract (NTS) (Extended Data Fig. 1b). However, these HFD-induced postprandial morphological changes in microglia were restrained to some discrete brain areas, and were not detected in the dorsomedial hypothalamus (DMH), the dorsal motor nucleus of the vagus (DMV), the CA3 area of the hippocampus nor in the S1BF cortical area (Extended Data Fig. 1c,d). These morphological changes were not detected in the ARC at 1 hour after HFD introduction (Extended Data Fig. 1e), and were totally reversed after 1 day, i.e. in mice fed with SD for 21 hours after the 3h-HFD exposure (Extended Data Fig. 1f). HFD-induced postprandial microglial hyper-ramification in the ARC was not found in female mice (Extended Data Fig. 1g).

**Figure 1.**
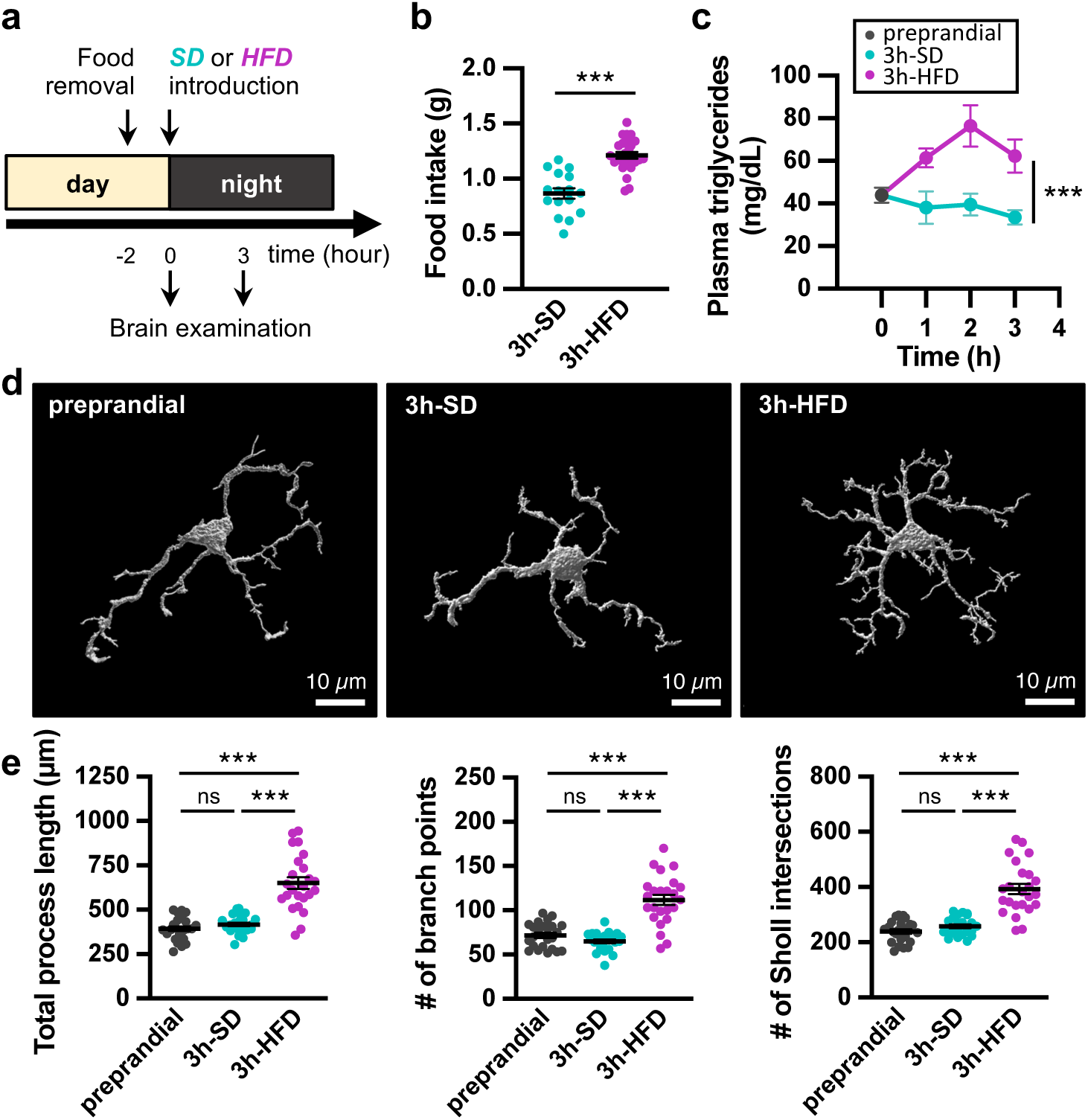
HFD consumption for 3 hours induces hyper-ramification of microglia in the arcuate nucleus of the hypothalamus in mice. **(a)** Timeline and experimental groups to investigate prandial states in 8/9-week-old male C57BL/6J mice. Food was removed for 2 hours at the end of the inactive light phase, and then standard diet (SD) or high fat diet (HFD) was introduced at light off for 3 hours. Mice were analysed before (preprandial state) or after feeding (postprandial states). **(b)** Food intake of mice fed for 3 hours with either standard (3h-SD) or high fat diet (3h-HFD) (3h-SD: n= 16 mice; 3h-HFD: n= 26 mice). **(c)** Plasma triglycerides excursion over a 3-hour-period of feeding with either SD or HFD (preprandial, n=23 mice; 3h-SD: n= 6 mice for each time; 3h-HFD: n= 5-6 mice for each time point). **(d)** Representative 3D reconstructed images of microglia in the arcuate nucleus of the hypothalamus of mice in the preprandial state or in the postprandial state, upon standard (3h-SD) or high fat diet (3h-HFD). Microglia were immunodetected in fixed brain slices using anti-Iba1 antibodies and imaged by confocal microscopy. Iba1^+^ cells were reconstructed in 3-dimensions using Imaris software. **(e)** Morphometric characteristics of arcuate Iba1^+^ cells according to the prandial state (preprandial, n=24 cells from 4 mice; 3h-SD, n=24 cells from 4 mice; 3h-HFD, n=24 cells from 4 mice). Automatic quantification of cell morphology was done using Imaris software. **Statistics.** Data are mean ± standard error of the mean (SEM). Asterisks indicate statistical differences between groups: ***p<0.001; ns, non significative. Data were analysed using two-tailed unpaired Student’s test (in b), two-way ANOVA (in c), and one-way ANOVA and Tukey’s tests (in e).

Quantification of Iba1^+^ cells on brain sections revealed no increase in microglia density in the hypothalamus after 3h-HFD neither in the ARC, the ventromedial hypothalamus (VMH) nor in the DMH, compared with the preprandial situation (Extended Data Fig. 1h,i). FACS analysis of freshly dissociated cells from mediobasal hypothalamus showed no change in the fraction of CD11b^+^-CD45^low^-CD206^-^-P2RY12^high^ cells regardless of the prandial state, confirming the stability of the microglia density in this area after 3h-HFD (Extended Data Fig. 1j,k,l). In addition, we did not observe any change in the proportion of lymphoid cells, macrophages, or granulocytes in the mediobasal hypothalamus after 3h-HFD, indicating the absence of peripheral immune cells infiltration during this postprandial period. Finally, we did not detect any variation in the expression of CD45, CD206 and P2RY12 markers in hypothalamic microglia and macrophages (Extended Data Fig. 1m). Altogether, these results reveal that postprandial conditions can trigger a transient reactive state in hypothalamic microglia, but without overt gliosis. This acute and moderate postprandial response is dependent on the composition of the food ingested, occurring after consumption of HFD but not following SD.

### The microglial response in the hypothalamus is induced by the postprandial hypertriglyceridemia

We next investigated the biological mechanisms underlying the HFD-dependent acute reaction of hypothalamic microglia. Pair-feeding experiments demonstrated that the postprandial microglial response was dictated by the nature of ingested food rather than the quantity consumed per se (Extended Data Fig. 2a). In addition, the acute HFD-induced microglial response developed when triglyceridemia increased (Fig. 1c). Therefore, nutritional lipids entering the bloodstream represent prime candidates for inducing the postprandial microglial response. Consistent with this hypothesis, previous studies reported rapid uptake of blood triglyceride-rich lipoparticles by hypothalamic microglia ^14^. Thus, we designed experiments to investigate the relation between postprandial hypertriglyceridemia and acute microglial reactivity in C57BL/6J male mice. We first substituted HFD by standardized oral gavage with peanut oil, while water was given as control (Fig. 2a). As expected, peanut oil gavage triggered hypertriglyceridemia (Fig. 2b). It also caused microglial hyper-ramification after 3 hours (Fig. 2c,d). Next, we tested whether systemic administration of lipids bypassing the gastrointestinal tract could still trigger the microglial response. The lipid emulsion of Olimel, a solution used for parenteral nutrition, was injected intraperitoneally in mice, while saline was administered to controls (Fig. 2e). Olimel injection induced a strong and transient hypertriglyceridemia (Fig. 2f), and triggered pronounced microglial hyper-ramification 3 hours after injection (Fig. 2g,h). Notably, intraperitoneal injection of glucose (0.5 g/kg) mimicking postprandial hyperglycemia ^31^ did not affect microglial morphology (Extended Data Fig. 2b). These data demonstrate that blood-borne nutritional lipids are sufficient to trigger the acute HFD-stimulated postprandial microglial response.

**Figure 2.**
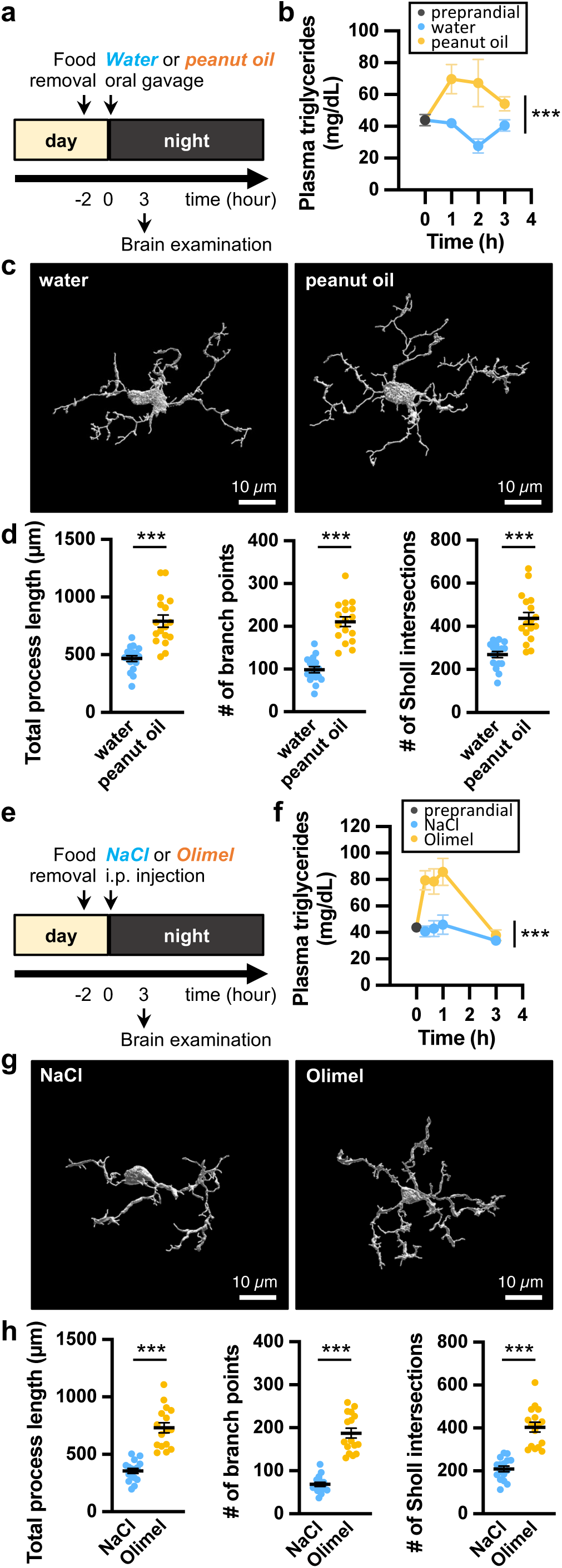
Postprandial hypertriglyceridemia triggers acute hyper-ramification of microglia in the arcuate nucleus of the hypothalamus in mice. **(a)** Experiment 1 designed to investigate the relation between acute hypertriglyceridemia and microglial reactivity in the arcuate nucleus of the hypothalamus. Hypertriglyceridemia was induced by a gavage with peanut oil (200 µl). Peanut oil or water (as a control) were given at light off. Brains were collected 3 hours after gavage. **(b)** Plasma triglycerides excursion after peanut oil or water gavage in 9/11-week-old C57BL/6J male mice (n=6 mice for each time). **(c)** Representative 3D reconstructed images of microglia in the arcuate nucleus of the hypothalamus of mice force-fed with peanut oil or water. Microglia were immunodetected in fixed brain slices using anti-Iba1 antibodies and imaged by confocal microscopy. Iba1^+^ cells were reconstructed in 3-dimensions using Imaris software. **(d)** Morphometric characteristics of arcuate Iba1^+^ cells after peanut oil or water gavage (Water, n=17 cells from 4 mice; Peanut oil, n=17 cells from 4 mice). Automatic quantification was done using Imaris software. **(e)** Experiment 2 designed to investigate the relation between acute hypertriglyceridemia and microglial reactivity in the hypothalamus. Hypertriglyceridemia was induced by intraperitoneal injection of the fat emulsion of Olimel (16 µl/g). Olimel or NaCl (as a control) were given at light off. Brains were collected 3 hours after injection. **(f)** Plasma triglycerides excursion after Olimel or NaCl ip injection in 10/14-week-old C57BL/6J male mice (n=5-6 mice for each time). **(g)** Representative 3D reconstructed images of microglia in the arcuate nucleus of the hypothalamus of mice injected with Olimel or NaCl. Microglia were immunodetected in fixed brain slices using anti-Iba1 antibodies and imaged by confocal microscopy. Iba1^+^ cells were reconstructed in 3-dimensions using Imaris software. Scale bar, 10 µm. **(h)** Morphometric characteristics of arcuate Iba1^+^ cells after Olimel or NaCl injection (NaCl, n=16 cells from 4 mice; Olimel, n=16 cells from 4 mice). Automatic quantification was done using Imaris software. **Statistics.** Data are mean ± standard error of the mean (SEM). Asterisks indicate statistical differences between groups: ***p<0.001. Data were analysed using two-tailed Mann-Whitney test (in d, h), and two-way ANOVA (in b, f).

### Consumption of high-fat diet for 3 hours reconfigures the translatome of mediobasal hypothalamic microglia

The morphological response of microglia to short-term HFD is remarkably rapid. To identify the associated molecular response and capture immediate, protein-synthesis-dependent changes reflected by mRNAs actively engaged in translation within microglia, we implemented the RiboTag strategy upon Cre-dependent recombination^32^. This approach provides a comprehensive and accurate analysis of the microglial translatome^33^. In brief, *Cx3cr1*^CreER/+^:Rpl22^HA/+^ mice, hereafter referred to as RiboTag^Mg^ mice, were generated and used to isolate ribosome-associated mRNA from mediobasal hypothalamic microglia (Extended Data Fig. 3a,b,h). We first demonstrated that RiboTag^Mg^ mice efficiently expressed the HA tag in hypothalamic Iba1-positive microglia (Extended Data Fig. 3c), exhibited normal eating behavior and HFD-induced postprandial hypertriglyceridemia (Extended Data Fig. 3d,e) and presented typical HFD-induced postprandial microglial response (Extended Data Fig. 3f,g). RNA sequencing of actively translated mRNAs, identified 14,212 expressed genes, with a very significant enrichment in microglia markers compared to other brain cells markers (supp-Fig 3i). To isolate genes associated with the acute HFD-mediated postprandial microglial response, we applied Bayesian clustering and ranked genes according to their expression profiles across the prandial states (Extended Data Fig. 3j). Genes were then assigned to one of four predefined transcriptional models: a *flat* model containing 7856 unresponsive genes, a *time and food interaction* model containing 3854 genes responding to 3 hour-food intake independently of the diet composition, a standard diet specific model containing 351 responsive genes regulated by the 3h-SD, and a fat intake-correlated model containing 2,151 responsive genes regulated by the 3h-HFD of which 1,134 were downregulated and 1,017 upregulated (Fig. 3a; Extended Data Fig. 3k,l). Because of the specific morphological effect of HFD-mediated postprandial microglial response, we then focused on fat intake-correlated genes. Principal component analysis (PCA) using the 2,151 genes showed that samples from a particular prandial state segregated together (Fig. 3b), indicating that nutrient intake triggered distinct transcriptomic responses in hypothalamic microglia. Out of the 2,151 genes, we identified 894 genes that explained most of the variation among the 3 experimental groups (dimension 1; 52.2%) (Fig. 3c). Plotting these genes on a heatmap highlighted the changes in the translatome landscape in microglia following fat intake (Fig. 3d). Computational enrichment analyses on annotated genes (GSEA) using the Gene-Ontology Biological Process database revealed that fat intake affected a variety of biological processes with network analysis showing that these latter could be grouped in nine main processes (Fig. 3e). Interestingly, genes positively correlated with fat intake promoted synaptic signaling, synapse assembly and cell development, while those negatively correlated were associated with homeostatic, biosynthetic, and metabolic processes, cell cycle, intracellular transport, and immune function. Consistent results were obtained using the Gene-Ontology Cellular Component database, with genes positively correlated with fat intake being preferentially localized in synapses and dendrites while those negatively correlated with fat intake were preferentially localized in vacuoles and organelles (Extended Data Fig. 3m). Top upregulated genes (i.e. *Dok7, Atpsckmt, Shisa9, Lrrtm4, Myrf, Hrh1, Lingo2, Adgrb3, Grm8, Grid2, Grm7, Syt6*) are known to be involved in synapses organization and stabilization, and in histaminergic and glutamatergic signaling. On the contrary, top downregulated genes (i.e. *Coro7, Npnt, Pecam1, Parn, Arsg, Eya4, Card11, Akip1, Chlsn, C1rl, Fancd2, Sh2d6*) are known to be involved in vesicle trafficking and degradation of macromolecules (Fig. 3f,g; Extended Data Fig. 3n). Of note, our analysis also revealed that the fat-intake postprandial transcriptomic response consisted in deregulation of a large number of unannotated genes whose function is currently unknown (Extended Data Fig. 3o). As a whole, these data reveal a fast fat-dependent molecular response in microglia from the mediobasal hypothalamus within 3 hours of a meal. Functional analyses of the fat-regulated genes suggest that these responses consist in a switch from immune surveillance and phagocytic activity to neuromodulation of synapses.

**Figure 3.**
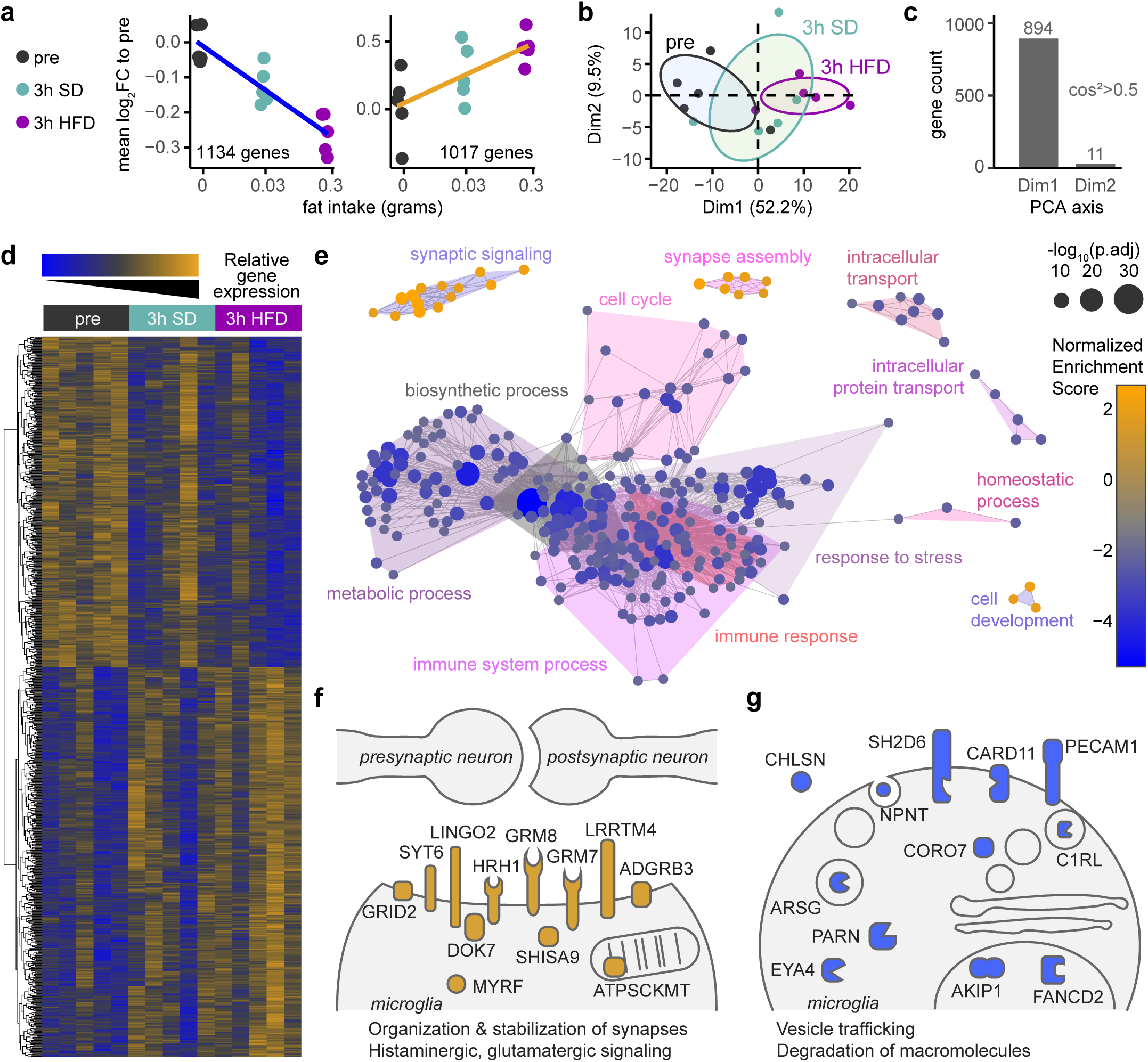
HFD-induced postprandial reactive microglia. **(a)** Representation of RiboTag data from microglia. Data is mean log2 fold-change to the pre-prandial group of RiboTag data across all identified genes for each individual. SEM is too small to be visible. The blue line represents a negative linear correlation to the log_10_ fat intake, whereas the orange line represents a positive correlation. **(b)** PCA on genes correlated to the fat intake. The dimension 1 of the PCA separates the individuals by the time and the quantity of fat intake. **(c)** Barplot of the number of genes identified as associated to dimension 1 with a cos²>0.5. Only 11 of these genes are also correlated to the dimension 2. **(d)** Heatmap of genes associated to dimension 1 from PCA. RiboTag data are centered and scaled. Individuals are sorted by their lipid intake. **(e)** Network of GSEA results of Gene Ontology Biological Processes. A link was drawn between two categories if a smaller category shares 90% homology in its leading edges genes composition with a larger category. The size represents the -log_10_ adjusted p value, and the color the normalized enrichment score. The most central category is used to identify each area of the network.

### Genetic ablation of *Asc* in microglia inhibits the postprandial HFD-dependent microglial response

To assess the behavioral outcome of the postprandial microglial response, we sought to interfere with the microglial activity. In the nervous system, the Apoptosis-associated Speck-like protein containing a CARD (ASC) is mainly expressed in microglia (Extended Data Fig. 4a,b), where it acts as a master regulator controlling metabolism, phagocytosis and inflammatory responses ^34–37^. ASC also regulates cytoskeleton rearrangement and cell morphology ^38^. Thus, to alter the postprandial microglial response to short term HFD, we selectively deleted the *Asc* gene in adult microglial cells. Mice carrying conditional floxed *Asc* alleles (*Asc*^fl/fl^ mice ^39^) were crossed with mice expressing the tamoxifen-inducible Cre recombinase under the direction of the *Cx3cr1* promoter (*Cx3Cr1*^CreER/CreER^ mice) ^40^ (Extended Data Fig. 5a). Tamoxifen treatment was given to the *Cx3cr1*^CreER/+^-*Asc*^fl/fl^ progeny to produce Cre-mediated deletion of *Asc* gene in microglia (Extended Data Fig. 5b). Tamoxifen-injected mice with *Asc* knock-out in microglia were named *Asc*^MgKO^ mice. Control mice were *Cx3cr1*^CreER/+^ harboring wild-type *Asc* alleles. Both control and KO mice received tamoxifen and were studied 5-6 weeks after treatment. To validate *Asc* gene deletion, we performed a series of molecular and functional analyses. *Asc* deletion in genomic DNA of FACS-sorted CD11b^+^-CD45^low^ microglia, CD11b^-^ non-microglial brain cells, and Ly6G^-^-CD11b^+^ blood monocytes from control and *Asc*^MgKO^ mice was evaluated by PCR (Extended Data Fig. 5c-f). 30 days after tamoxifen treatment, *Asc* deletion was found in microglia, but not in non-microglial cells, or blood monocytes (Extended Data Fig. 5c-g). We confirmed that Cre-mediated recombination was stable over time in long-lived CX3CR1^+^ microglia (up to 120 days post-tamoxifen treatment), but transient in short-lived CX3CR1^+^ cells, such as monocytes, which are rapidly replaced throughout continuous differentiation of unrecombined bone-marrow-derived progenitors (Extended Data Fig. 5g) ^41^. No deletion can be found in control mice (Extended Data Fig. 5h). Using real-time RT-qPCR on total mRNA from isolated cells, we found that *Asc*^MgKO^ exhibited a drastic decrease by 96% in microglial *Asc* mRNA expression while no change could be noticed in non-microglial cells in the brain, in peritoneal macrophages and in bone-marrow-derived macrophages (Fig. 4a). Finally, primary microglial cultures sorted from newborn mice, with quite complete *Asc* gene deletion and large ASC protein deficiency induced in vitro by 4-hydroxytamoxifen (Extended Data Fig. 6a-c), failed to assemble ASC specks when stimulated by LPS and nigericin (Fig. 4b; Extended Data Fig. 6c). Altogether, these data show the specific ablation of *Asc* gene in microglia along with altered ASC functioning in these cells in *Asc*^MgKO^ mice.

**Figure 4.**
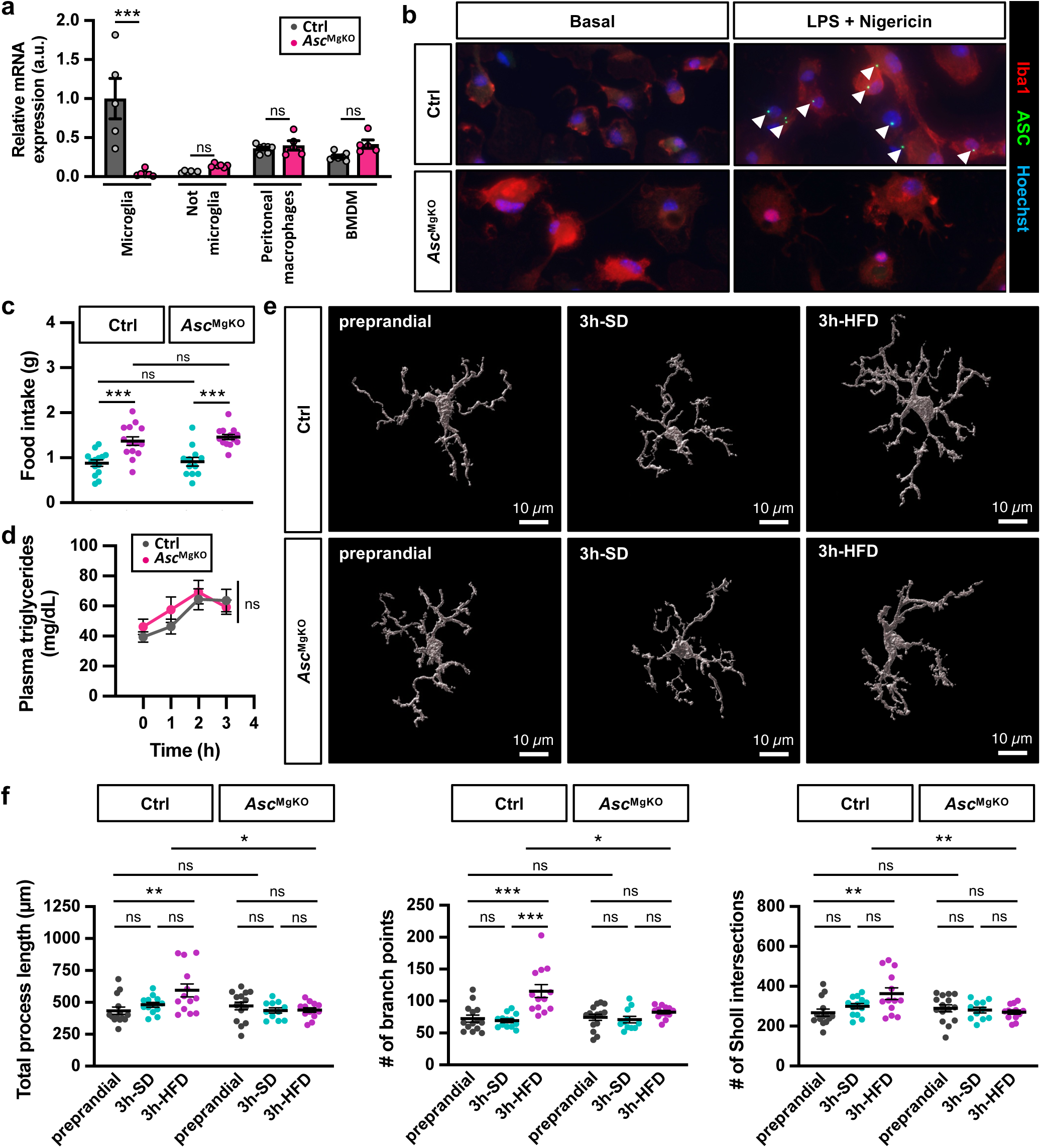
*Asc* gene deletion in microglia alters HFD-induced postprandial microglial reactivity. (a) Relative expression of *Asc* mRNA in FACS-sorted microglial cells (CD11b^+^/CD45^low^) and not microglial cells (CD11b^-^) from the brain of Cx3cr1^Cre/+^-*Asc*^+/+^ (Ctrl) and Cx3cr1^Cre/+^-*Asc*^fl/fl^ (*Asc*^MgKO^) male mice, 30 days after tamoxifen injections. Relative expression of *Asc* mRNA in primary peritoneal macrophages and BMDM were also analysed in both genotypes, at the same experimental timing. All mRNA levels were quantified relative to Polr2a housekeeping gene expression by the ΔΔCT method and presented as fold-change relative to the Ctrl group, whose mRNA level was standardized to 1 (Ctrl: n= 5-7 mice; *Asc*^MgKO^: n= 5-7 mice). **(b)** Immunodetection of ASC specks in LPS-primed nigericin-treated Ctrl and *Asc*^MgKO^ microglia. Cells were cultured from newborn Cx3cr1^Cre/+^-*Asc*^+/+^ (Ctrl) and Cx3cr1^Cre/+^-*Asc*^fl/fl^ mice and were all treated with 4-hydroxytamoxifen in vitro. After ASC depletion, cells were stimulated with LPS and nigericin, fixed with paraformaldehyde, and stained with anti-ASC antibody (green), anti-Iba1 antibody (red), and Hoechst dye (blue). Arrowheads show immunolabeled green fluorescent ASC specks. **(c)** Food intake of Ctrl and *Asc*^MgKO^ male mice fed for 3 hours with either standard (3h-SD) or high fat diet (3h-HFD) (Ctrl: n= 14 mice; *Asc*^MgKO^: n= 12-14 mice). **(d)** Plasma triglycerides excursion in Ctrl and *Asc*^MgKO^ male mice over a 3-hour-period of feeding with either SD or HFD (Ctrl: n= 10-12 mice; *Asc*^MgKO^: n= 9-11 mice). **(e)** Representative 3D reconstructed images of microglia in the arcuate nucleus of the hypothalamus of Ctrl and *Asc*^MgKO^ male mice in the preprandial state or in the postprandial state, upon standard (3h-SD) or high fat diet (3h-HFD). Microglia were immunodetected in fixed brain slices using anti-Iba1 antibodies and imaged by confocal microscopy. Iba1^+^ cells were reconstructed in 3-dimensions using Imaris software. **(f)** Morphometric characteristics of arcuate Iba1^+^ cells in Ctrl and *Asc*^MgKO^ male mice according to the prandial state (Ctrl: preprandial, n=14 cells from 3 mice; 3h-SD, n=13 cells from 3 mice; 3h-HFD, n=13 cells from 3 mice; *Asc*^MgKO^: preprandial, n=15 cells from 3 mice; 3h-SD, n=11 cells from 3 mice; 3h-HFD, n=13 cells from 3 mice). Automatic quantification of cell morphology was done using Imaris software. **Statistics.** Data are mean ± standard error of the mean (SEM). Asterisks indicate statistical differences between groups: **p<0.01; ***p<0.001; ns, non significative. Data were analysed using Kruskal-Wallis and Dunn’s tests (in a, f), two-way ANOVA and Sidak tests (in c), and mixed-effects model’s tests (in d).

*Asc*^MgKO^ mice did not express obvious behavioral anomalies or deficits. Phenotyping and gross necropsy revealed comparable growth, size and organ weights between *Asc*^MgKO^ and control mice (Extended Data Fig. 7a,b). Plasma chemistry and complete blood count did not show diabetes, anemia, leukemia, infection, or immune system abnormalities in *Asc*^MgKO^ mice compared to control mice (Extended Data Fig. 7c-e). Post-mortem brain examination did not show any malformation in *Asc*^MgKO^ mice and histological analyses reported comparable anatomy, myelination, neuron and microglia densities in the hypothalamus of *Asc*^MgKO^ and control mice (Extended Data Fig. 7f-i).

We then examined postprandial responses in male *Asc*^MgKO^ and control mice. Both groups exhibited normal food intake on SD or HFD for 3 hours, and developed comparable hypertriglyceridemia within 3 hours following HFD introduction (Fig. 4c,d). Microglia morphology in the ARC was also similar between genotypes in the preprandial state, mirroring that of C57BL/6J mice (Fig. 4e,f). As in C57BL/6J mice, 3h-SD did not alter the microglial morphology in either *Asc*^MgKO^ or control mice. In contrast, whereas 3h-HFD triggered the typical postprandial microglial hyper-ramification in control mice, this response was completely abolished in *Asc*^MgKO^ mice. These findings indicate that ASC in microglia does not alter microglial morphology under resting conditions or following a standard meal, but is essential for the HFD-induced postprandial morphological microglial response.

### Genetic ablation of *Asc* in microglia increases consumption of dietary fat

We next studied the feeding behavior of *Asc*^MgKO^ and control male mice across several days. Ad libitum food intake was recorded over a 72-hour period. The analysis revealed that ASC deficiency in microglia had no consequence on food intake when mice were fed with a SD (Fig. 5a,b), but significantly increased the cumulative food intake on HFD. It also changed the temporal organization of the feeding behavior. Although the amount of food ingested per bout and the duration of bout remained unchanged, the number of feeding bouts was strongly increased in *Asc*^MgKO^ mice kept on HFD. These effects developed progressively over time, becoming significant after 15 hours and maximal after 2 days (Fig. 5c). HFD-associated overfeeding in *Asc*^MgKO^ mice was maintained and even exacerbated after a 21-hour fast (Fig. 5d). Experiments in female *Asc*^MgKO^ mice showed that HFD-associated overfeeding developed in males only (Extended Data Fig. 8a,b). These results indicate that ASC in microglia has no effect on food intake during normal physiology when eating a SD, but is required to promote satiety and limit overfeeding on HFD. This HFD-dependent anorexigenic mechanism takes place in male but not in female mice.

**Figure 5.**
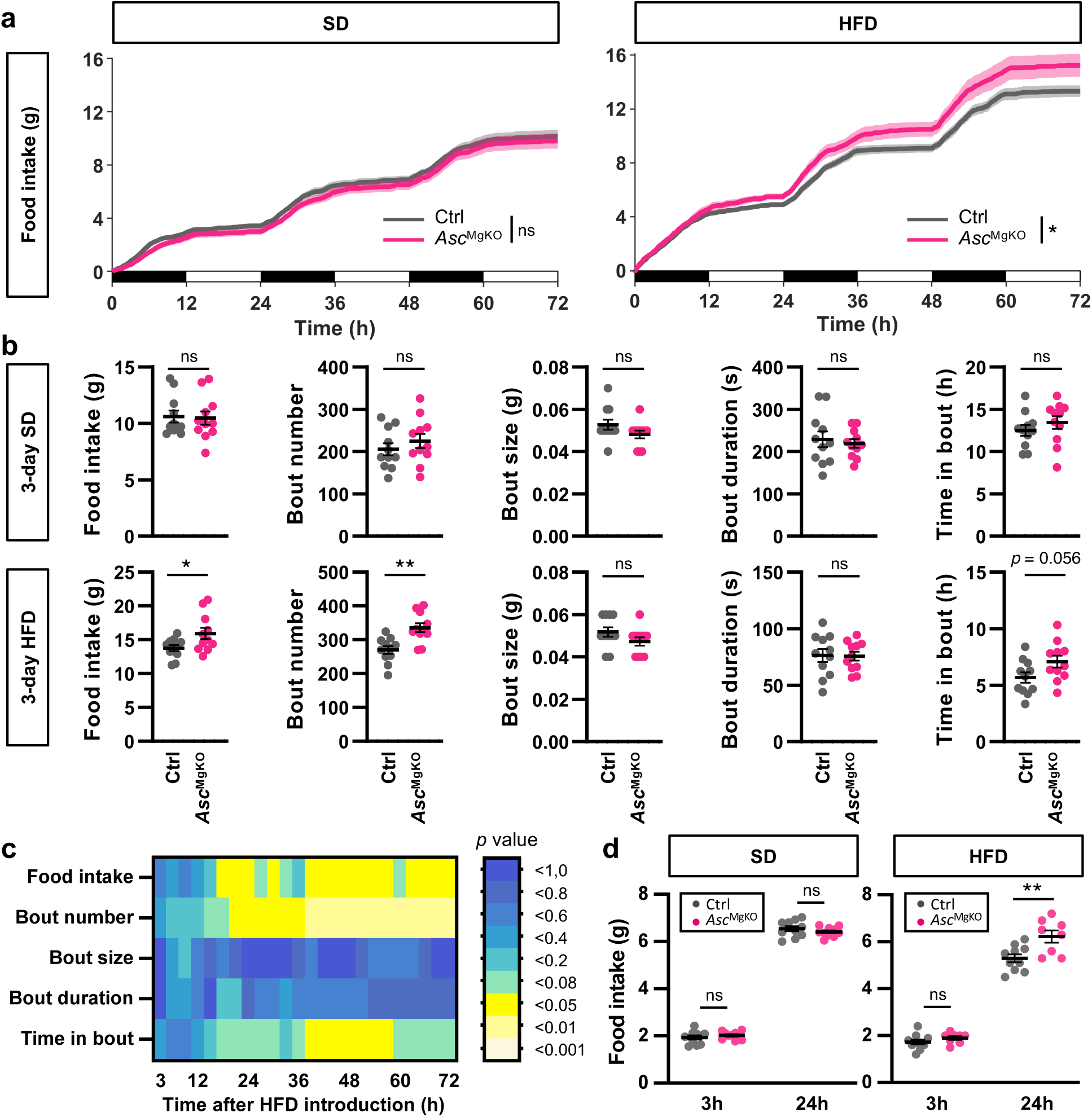
*Asc* gene deletion in microglia raises overfeeding in male mice when consuming HFD. **(a)** Cumulative food intake of 12-week-old male Ctrl and *Asc*^MgKO^ mice fed with SD or HFD ad libitum for 72 hours (Ctrl: n=11 mice; *Asc*^MgKO^: n=11 mice). **(b)** Microstructure of food intake including total food intake, number of feeding bouts, mean size of feeding bouts, mean duration of feeding bouts, and percentage of time spent in feeding bouts in 12-week-old male Ctrl and *Asc*^MgKO^ mice fed with SD or HFD ad libitum for 72 hours (Ctrl: n=11 mice; *Asc*^MgKO^: n=11 mice). **(c)** Heatmap showing statistical differences in the microstructure of food intake. The scale bar represents the color-coded p values which were calculated for every 3h period. **(d)** Cumulative food intake of 9/12-week-old male Ctrl and *Asc*^MgKO^ mice fed with SD or HFD ad libitum for 24 hours after a 21-hour-fast. Refeeding was measured manually after 3h or 24h (Ctrl: n=10 mice; *Asc*^MgKO^: n=8-10 mice). **Statistics.** Data are mean ± standard error of the mean (SEM). Asterisks indicate statistical differences between groups: *p<0.05; **p<0.01; ***p<0.001; ns, non significative. Data were analysed using two-way ANOVA (in a), and two-tailed unpaired Student’s test (in b, c, d).

### The postprandial HFD-dependent microglial response HFD consumption is lost in obese mice

The HFD-dependent postprandial microglial response was characterized in young adult, healthy male mice. We next asked whether this physiological mechanism was preserved in the context of obesity. Thus, we compared microglial morphology in the ARC of age-matched male lean C57BL/6J, and obese Lep^KO^ (ob/ob) and Mc4r^KO^ mice (Fig. 6a). Each strain was studied at preprandial and 3h-HFD postprandial state. Notably, compared to lean mice, all strains of obese mice were hyperphagic (Fig. 6b) and exhibited hypertriglyceridemia 3 hours after HFD introduction (Fig. 6c). We also observed some subtle differences in the preprandial state, with microglia from obese mice exhibiting higher branching compared to lean control (Fig. 6d,e). Strikingly, whereas 3h-HFD induced the expected extension and ramification of microglial processes in lean mice, no morphological changes were detected in obese mice following 3h-HFD. These findings indicate that the postprandial HFD-induced microglial response is lost in obese mice despite a higher fat load.

**Figure 6.**
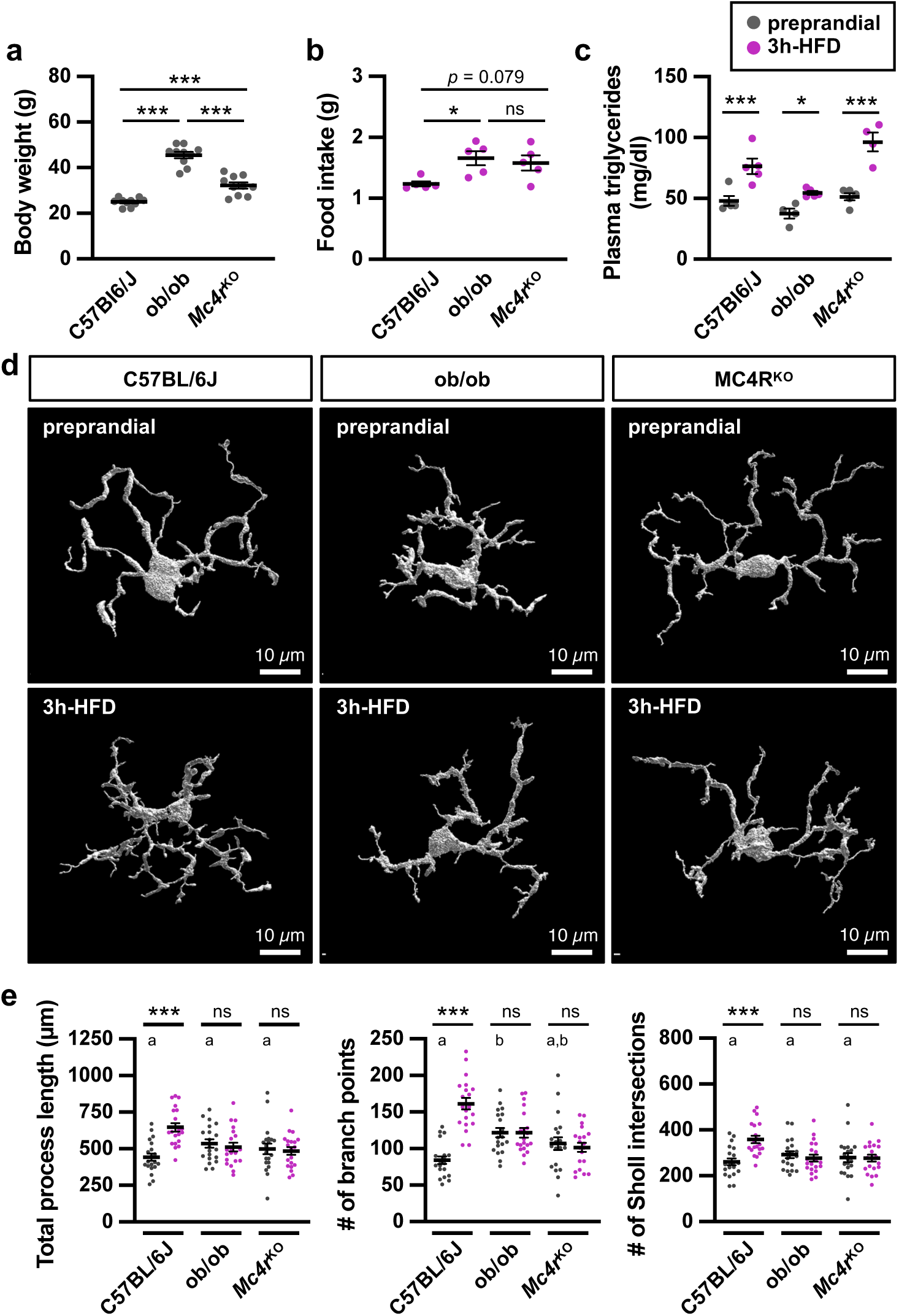
Postprandial hypertriglyceridemia does not induce microglial reactivity in obese mice. Comparison of the HFD-induced acute microglial response in 8/10-week-old C57BL/6J, ob/ob, Mc4r^KO^ male mice. **(a)** Body weight of the mice (n=10 per genotype). **(b)** Food intake of the mice fed for 3 hours with high fat diet (3h-HFD) (n=5 per genotype). **(c)** Plasma triglycerides levels in C57BL/6J, ob/ob, Mc4r^KO^ male mice after a 3-hour-period of feeding with HFD (n=4-5 per genotype). **(d)** Representative 3D reconstructed images of microglia in the arcuate nucleus of the hypothalamus of 8/10-week-old C57BL/6J, ob/ob, Mc4r^KO^ male mice in the preprandial state or fed a high-fat diet for 3 hours (3h-HFD). Microglia were immunodetected in fixed brain slices using anti-Iba1 antibodies and imaged by confocal microscopy. Iba1^+^ cells were reconstructed in 3-dimensions using Imaris software. **(e)** Morphometric characteristics of arcuate Iba1^+^ cells in lean and obese mice (C57BL/6J: preprandial, n=20 cells from 4 mice; 3h-HFD, n=20 cells from 4 mice; ob/ob: preprandial, n=20 cells from 4 mice; 3h-HFD, n=20 cells from 4 mice; Mc4r^KO^: preprandial, n=20 cells from 3 mice; 3h-HFD, n=20 cells from 4 mice). Automatic quantification was done using Imaris software. **Statistics.** Data are mean ± standard error of the mean (SEM). Asterisks indicate statistical differences between groups: *p<0.05; ***p<0.001; ns, non significative. For multiple comparisons, different letters above the bars indicate statistically significant difference (p<0.05). Data were analysed using one-way ANOVA and Tukey’s tests (in a, b, c, e), and two-way ANOVA and Tukey’s tests (in e).

## Discussion

This study demonstrates that microglia in the mouse brain respond acutely to ingested foods at the time scale of a meal, undergoing morphological remodeling that reflect functional adaptation. Transcriptomics data indicate that following a fatty meal, microglia temporarily shift their roles related to innate immunity and brain surveillance toward modulation of neurotransmission. Selective inhibition of ASC-related microglial functions impaired postprandial microglial reactivity and altered feeding behavior, underscoring the critical role of microglial responses in animal physiology, particularly in the brain control of food intake.

Detailed histological analysis reveals hyper-ramification of microglia after a meal. This finding is consistent with our previous work showing a significant increase in cell membrane capacitance in microglia few hours after food introduction ^28^. In addition, we found that the postprandial microglial response is macronutrient-dependent, being specifically induced by high-fat food. This effect seems independent from sensory cues, other post-ingestive signals, or endogenous rhythmicity, since standard food, glucose or water intake did not induce it. Rather, artificial elevation of lipids in the blood was sufficient to stimulate the HFD-associated microglial response. Whether lipid sensing by the microglia is direct remains to be elucidated. Yet, microglia express a variety of molecular sensors of nutritional lipids on their surface, including LPL, TREM2, TLR4, and CD36 ^23,42–44^. These receptors can either hydrolyze triglycerides, sense saturated fatty acids and/or are able to promote the uptake of circulating endogenous lipoproteins or free fatty acids ^14,42^. Consistent with this, we found that high fat diet (7% butter, 7% lard, coconut oil, soja oil), peanut oil (100%) and lipid emulsion of Olimel® (80% olive oil, 20% soja oil), which all contain saturated fatty acids, induced microglia hyper-ramification in a similar manner. Additional nutritional studies could identify the lipidic determinants of the postprandial microglial response.

Postprandial reactive microglia were not disseminated through the entire brain but were anatomically confined to the ARC of the hypothalamus and the NTS of the brainstem. These regions constitute a distinct microenvironment flooded with blood-borne molecules, including hormones and nutrients, that diffuse from fenestrated capillaries ^8^. This anatomical characteristic may account for the regioselectivity of the microglial response to dietary lipids. Furthermore, previous transcriptional profiling has identified distinct microglial sub-populations that are neuroanatomically segregated ^45^. It is therefore plausible that microglia in the ARC and NTS possess specific molecular machinery that enhances their responsiveness to lipids. Additionally, our findings indicate that the postprandial microglial response does not occur in females. This sexual dimorphism may be associated with molecular and anatomical variations influenced by steroid hormones ^46–48^.

Transcriptomics data reveal the rapid reprogramming of gene expression in microglia located in the mediobasal hypothalamus within 3 hours of HFD exposure. This suggest that reactive microglia shifted from immune surveillance and phagocytosis to synaptic modulation. In particular, HFD-induced genes in microglia such as *Adgrb3*, *Atpsckmt*, *Dok7, Lingo2, Syt6* modulate synaptic strength and remodeling, and others such as *Hrh1, Grm7, Grm8, Grid2, Lrrtm4* and *Shisa9* modulate histaminergic and glutamatergic signaling. Concurrently, FACS analysis and histological examination did not show any increase in Iba1-immunoreactivity, upregulation of inflammatory markers, or changes in microglia cell density or white blood cells infiltration. Collectively, these findings suggest that a 3 hour-HFD exposure does not elicit a robust inflammatory microglial response, contrasting with the more pronounced effects reported after days or months of HFD exposure ^9,13–15,19,49,50^. These results align with others studies describing minimal microgliosis, if any, during very short term HFD ^49,51,52^. Furthermore, our research reveals that microglia employ an ASC-dependent mechanism to undergo morphological changes and modulate feeding behavior. Thus, microglia mobilize a conserved signaling pathway involved in innate immunity and inflammation to operate physiological functions. Finally, we identified a unique microglial plasticity following a fatty meal, characterized by hyper-ramification, enhanced neuromodulatory activity, and bodily functions.

Reactive microglia have a complex impact on brain function, potentially serving as sources of neurotoxic agents that drive neurological diseases ^53^. Chronic consumption of a hypercaloric diet activates microglia in the hypothalamus, contributing to obesity ^13–16,19^. Here we found that genetic deletion of microglial *Asc* significantly alters feeding behavior, leading to increased consumption during the initial days of HFD. This indicates that the acute microglial response to short term HFD has a positive adaptive value, normalizing food intake during periods of overfeeding. Notably, reactive astrocytes have been shown to play a similar homeostatic role in response to acute high-fat feeding ^54^. These acute glial reactions complement a set of anorectic responses to positive energy balance deviations, including changes in neuronal sensitivity, gene expression, synaptic organization and cell renewal within brain circuits that regulate food intake ^51,55–57^. During short term HFD, microglia are also essential for the blood glucose regulation and multiple cognitive functions ^52^. In the long term, reactive microglia still display some beneficial effects, improving glucose tolerance and maintaining innate immunity ^23,24^.

Like the early morphological and transcriptomic response of microglia, regulation of feeding behavior by ASC is engaged on HFD only, consistent with a previous study showing that microglia are critical for optimal responses to dietary fat, and have no overt effect on baseline metabolism ^52^. It does not preclude the possibility of additional microglial functions in energy homeostasis when consuming a balanced diet. Notably, microglia facilitate the cephalic phase of insulin release on a standard diet, although this effect has only been observed after starvation ^27^. Fasting also influences the microglial control of glycemia ^52^. Here, the microglial-dependent regulation of food intake was more pronounced following after fasting. These findings suggest that metabolic functions of microglia are dynamic, probably limited in number after balanced meals, becoming more numerous and effective on the feeding behavior after drastic metabolic transitions, such as during fasting-refeeding cycles or when consuming high-fat high-calorie foods.

Genetic deletion of microglial ASC influences feeding behavior several hours after HFD introduction, emerging after the first signs of microglial remodeling. This delayed behavioral effect might be the accumulating result of a slow regulatory sequence between the microglial lipid sensing, the subsequent transcriptomic and morphological responses, and the terminal neuronal modulation. Previously, we described a similar delayed outcome following postprandial neuroglial plasticity after a standard meal, which relies at least in part on POMC activity ^31^. POMC neurons could also represent the final effectors of the microglial response. First, because microglia are reactive in the ARC, where POMC neurons are located, and second, because these neurons modulate feeding behavior with relatively slow kinetics ^58,59^. Although we examined several brain regions, we did not perform an exhaustive mapping of the postprandial microglial response throughout the brain and over the entire duration of HFD exposure. Thus, a response outside the ARC, potentially affecting other neuronal systems, cannot be ruled out. Notably, a microglial metabolic response, detectable 12 hours after HFD exposure, has been described beyond the hypothalamus, in the hippocampus and cortex ^52^, suggesting that brain microglia respond acutely to HFD in a spatiotemporal manner.

On the long term, restraining inflammatory signaling in microglia strongly limits diet-induced hyperphagia and weight gain ^19^. As in other chronic inflammatory diseases, transition from acute adaptive to sustained deleterious effects remains obscure. We found that microglial-dependent effects on feeding behavior upon HFD are maximal after 48 hours and then reduce. Others reported transient metabolic shift in microglia within first days on HFD ^52^. These data suggest that early microglial effects are temporary and might not be efficient on a long-term. Indeed, we found that the postprandial microglial response is lost in obese mice. Since the postprandial microglial response is anorectic, its absence can, to some extent, explain hyperphagia in obese animals.

In conclusion, here we characterized the dynamic postprandial microglial plasticity in the mouse brain, revealing that microglia rapidly respond to ingested food at the molecular and morphological levels. We demonstrated that microglia play a crucial role in heathy male mice, regulating food intake in a macronutrient-dependent manner. Our results also reveal that microglia employ an immune signaling to alert the brain and activate an anorectic response that curbs overeating during dietary fat consumption. As a whole, this study provides a novel regulatory mechanism of food intake, which may be overpassed in the context of obesity.

## Acknowledgments

We thank the animal facility of the CSGA in Dijon, especially Anne Lefranc, Martin Bertomeu, Laure Richardot, Patrick Cailler, and Virginie Cadiou for animal care. We thank the RAM-iExplore Facility in Montpellier for their help with the generation of the Ribotag mice. We also thank the animal facility of the Université Bourgogne Europe in Dijon, especially Valerie Saint-Giorgio, for help during rederivation service, and ethical considerations regarding animal experimentation. We thank the DImaCell imaging facility (Université Bourgogne Europe, Dijon), especially Elodie Noirot and Chrystel Deulvot for assistance. We thank the Imaflow facility (Université Bourgogne Europe, Dijon), especially Pierre Scaillierez, Audrey Geissler, and Amandine Bataille for histological studies, and Serge Monier, Nicolas Pernet, and Anabelle Sequeira Le Grand for FACS analysis. We thank Juliette Van Steenwinckel for her kind advices on the antibodies used for FACS, and François Hermetet for blood cell counting. We thank the MGX facility for the RNA sequencing, especially Camille Bellières and Hugues Parrinello. We thank students who assisted researchers, especially Savanah Buiret, Simon Cuenot, Sébastien Fargeton, Caroline Genevois, Laura Guenot, Audrey Mercier, Amélie Muller, Lucie Pagot, Rosana Pau Landa, Elsa Pélissier, Elise Pichler. We gratefully acknowledge Dr Mickaël Rialland and Dr Charles Thomas for their thoughtful inputs. This work was supported by Agence Nationale de la Recherche (contract MicroFlamEAT ANR-21-CE14-0033 to AB, VC, CR), by the French “Investissements d’Avenir” program/project ISITE-BFC (contract PlastEAT ANR-15-IDEX-0003 to AB), by the Région Bourgogne to AB, and by Fondation pour la Recherche Médicale (contract DEQ20170336744 to VP). We thank Vishva Dixit (Genentech) and the animal facility of the Université UCBL of Lyon (ANIRA-PBES, SFR Biosciences, Lyon), for providing *Asc*^KO^ and *Asc*^fl/fl^ mice. MGX facility acknowledges financial support from France Génomique National infrastructure, funded as part of “Investissement d’Avenir” program managed by Agence Nationale pour la Recherche (contract ANR-10-INBS-09).

## Methods

### Animals

C57BL/6J mice were purchased from Charles River. Transgenic mice, including *Cx3Cr1*^CreER^, *Rpl22*^HA^, Lep^KO^, and Mc4r^KO^ mice were obtained from Jackson Laboratories (Stock No. #020940, #029977, #000632, #032518, respectively). *Asc*^KO^ mice were obtained from Plateau de Biologie Expérimentale de la Souris (PBES facility, Lyon), kindly provided by Dr. Vishva Dixit (Genentech). *Asc* floxed mice from the Dr. Jurg Tschopp lab were kindly provided by Dr. Virginie Petrilli. *Asc*^MgKO^ mice were obtained by crossing *Asc*^fl/fl^ mice with *Cx3Cr1*^CreER/CreER^ mice. *Cx3cr1*^CreER/+^-*Asc*^fl/fl^ and *Cx3cr1*^CreER/+^-*Asc*^wt/wt^ progenies were used to produce *Asc*^MgKO^ and control mice, respectively, after tamoxifen treatment. Control and *Asc*^MgKO^ mice were both haploinsufficient for *Cx3cr1* (i.e Cre/+ heterozygous), on the same genetic background, and received tamoxifen equally. Floxed *Asc* alleles were carried by mutant mice only, while control mice kept wild type *Asc* alleles. This genetic scheme avoids misinterpretation due to the Cre expression itself or to the gene dosage of CX3CR1, especially given that microglial CX3CR1 signaling controls the degree of obesity^60^. RiboTag^Mg^ mice were generated by crossing tamoxifen-inducible *Cx3Cr1*^CreER/CreER^ mice ^40^ and *Rpl22*^HA/HA^ floxed mice ^32^. *Cx3Cr1*^CreER/+^-*Rpl22*^HA/+^ progeny was used to produce RiboTag^Mg^ mice after tamoxifen treatment. Animals were housed at 22.5 ± 1 °C on a 12h-12h reversed light-dark cycle (light off at 10h30; light on at 22h30). Cages contained enrichment and bedding material. Animals were acclimatized to their environment for at least 2 weeks before experiments. Before experiments, food (Cat. No A04, Safe) and water were given ad libitum. Experiments were made on 8/14-week-old mice. All protocols including animals were reviewed by our local ethic committee in strict accordance with European Community guidelines and agreed by the French Ministry of Higher Education and Research.

### Tamoxifen injection

To induce gene recombination in Cx3cr1^CreER/+^ mice, tamoxifen (Cat. No T5648, Sigma Aldrich) was dissolved in warm corn oil (Cat. No C8267, Sigma Aldrich) and administered by intraperitoneal injection (20 mg/ml; 75 mg/kg) once a day for 5 consecutive days at the age of 5 weeks. Mice were closely monitored during and after the treatment period. Experiments were carried out at least 30 days post tamoxifen injections.

### Experimental design

Mice were acclimatized for 3 days to single housing in a regular cage containing enrichment and bedding material. Mice were randomly assigned to 3 groups to compare different prandial states. All mice were submitted to a 2-hour food restriction just before lights off. At the beginning of the night period, mice belonging to the preprandial groups were killed. At this point, mice belonging to the postprandial groups received either a standard diet (SD; Cat. No A04, Safe) or a high-fat diet (HFD; Cat. No U8954v7, Safe). Postprandial mice were killed 3 hours after food reintroduction, or at another time described in the text. Food restriction standardized metabolic state and feeding behavior, ensuring substantial and reliable food intake after food reintroduction. As previously reported, the preprandial state is not a fasting state and does not alter level of metabolic hormones ^31^. To avoid stress, mice were trained for the 2-hour food restriction during 3 days. To prevent food neophobia, all mice received HFD pellets for 15 minutes the day before the experiment.

### Treatments

For oral lipid administration, mice received intragastric gavage with 200 µL of peanut oil (17% saturated fatty acids, mostly palmitic acid; Lesieur) or tap water as control. Gavage was given just before lights off. For direct systemic lipid administration, mice received an intraperitoneal injection of the lipid emulsion of Olimel® N7E (80% olive oil and 20% soy oil; Baxter) or NaCl 0.9% as control, just before lights off. Injected volume was dosed at 16 ml/kg.

### Automated food intake monitoring

Food intake was monitored in undisturbed mice using the BioDAQ System (Research Diets, Inc.). Mice were acclimatized for 3 days to single housing in a regular cage and feeding through a food hopper. Cages contained enrichment. Water was given ad libitum from regular bottles placed on the top of the cage. Mice showed normal food intake within 3 days of habituation. During habituation and experiments, food hoppers were closed for 2 hours before light off. For feeding behavior experiments, mice received SD for 3 days, and then HFD for 3 days. Data were collected continuously during experiments and were analyzed using the BioDAQ DataViewer software (Research Diets, Inc.). For analysis, the minimum food amount was filtered at 0.01 g per bout, and the maximum food amount at 0.2 g per bout.

### Fasting-refeeding study

Mice were acclimatized for 1 day to single housing in a regular cage, containing enrichment and bedding material. Water was given ad libitum from regular bottles placed on the top of the cage. Food was deprived for 21 hours. SD or HFD was then reintroduced and food intake was measured manually after 3 and 24 hours.

### Body composition

Lean mass and fat mass were analyzed by 3-in-1 Echo MRI Body Composition Analyzer (EchoMRI™).

### Blood analysis

Blood samples were collected in anesthetized mice by intracardiac puncture. For plasma triglycerides assay, blood was collected in heparin-containing tubes (Cat. No 41.1393.105, Sarstedt). For plasma insulin assay, blood was collected in EDTA-coated tubes (Cat. No 41.1504.005, Sarstedt). Samples were centrifuged at 4°C for 10 min at 2650 g. Plasma aliquots were stored at -80°C until use. Plasma triglycerides were measured using the colorimetric assay kit Triglycerides FS* (Cat. No 1 5710 00 10 026, DiaSys). The standard range was produced using a standard with a concentration of 200 mg/dL (Cat. No 1 5700 30 030, DiaSys). Plasma insulin was measured using AlphaLISA Insulin Immunoassay (Cat. No AL204C, Perkin Elmer). For complete blood count, blood was collected in EDTA-coated tubes (Cat. No 41.1504.005, Sarstedt). Cell blood count was performed on fresh samples using a hematology analyzer (scil Vet abc Plus+, Horiba). For blood glucose assay, blood was collected from the tail tip in conscious mice and blood glucose concentration was obtained with a glucometer (Accu-Chek® Compact Plus, Roche).

### Paraffin-embedded tissue histology

Mice were deeply anesthetized by intraperitoneal injection of a ketamine – xylazine mix (100 mg/kg – 10 mg/kg), and transcardiacally perfused with PBS (15 ml) and 4% paraformaldehyde prepared in 0.1 M phosphate buffer (40 ml). Brains were collected, post-fixed for 48 hours at 4°C, dehydrated in graded alcohols, cleared in xylene, and embedded in paraffin on Leica ASP300 Tissue processor. Slices of 5 µm thick were made on a Leica rotary microtome (Histocore Autocut R), and deposited on coated glass slides. Tissue sections were then deparaffinized in xylene, and rehydrated in decreasing concentrations of ethanol. For histological experiments, sections were stained with Cresyl Violet (Cat. No 5235, Merck) or Luxol Fast Blue (Cat. No 28106, Serva). Images of stained sections were realized on Axioscope A1 microscope (Zeiss) equipped with a Jenoptic Gryphax NAOS camera. For immunohistochemical experiments, rehydrated tissue sections were incubated in EDTA buffer pH 9 at 70°C for 2 hours to carry out antigen retrieving. Sections were further incubated in 3% hydrogen peroxide for 15 min, and in 3% bovine serum albumin for 20 min. Sections were then incubated for 1 hour at room temperature with primary antibody (rabbit anti-Iba1, 1:100; Cat. No CP290A, BioCare medical) and for 30 minutes at room temperature with a secondary antibody anti-Rabbit IgG conjugated with polymerase-HRP (ready-to-use ImmPress kit; Cat. No MP-7401, Vector Laboratories). Chromogenic revelation was performed using NovaRed kit (Cat. No SK-4800, Vector Laboratories). Sections were counterstained with Harris Hematoxylin (Cat. No 3801561E, Leica). After dehydration, tissue sections were mounted with organic mounting medium (Pertex; Cat. No LEIC811, Leica). Images of stained sections were captured with an Axio Imager microscope (Zeiss) equipped with an AxioCam camera (Zeiss).

### Frozen tissue histology

Mice were deeply anesthetized by intraperitoneal injection of a ketamine – xylazine mix (100 mg/kg – 10 mg/kg), and transcardiacally perfused with PBS (15 ml) and 4% paraformaldehyde prepared in 0.1 M phosphate buffer (40 ml). Brains were collected and post-fixed for 48 hours at 4°C. Tissues were then cryoprotected by immersion in 0.1 M phosphate buffer containing 30% sucrose, for 3 days at 4°C. Brains were included in OCT (Tissue-Tek® O.C.T. Compound; Cat. No 4583, Sakura), flash frozen in -65°C isopentane, and stored at -80°C until use. Serial 16-µm thick frontal sections were cut using a cryostat (CM3050S, Leica). Sections were mounted onto Superfrost Ultra Plus slides (ThermoScientific), and stored at -20°C until use. Sections were then processed for immunohistochemistry. Briefly, sections were incubated for 2 hours at room temperature in a blocking solution (NGS 1%, BSA 1%, Triton 0.3%, in PBS), and then incubated overnight at 4°C with a primary antibody prepared in the blocking solution (rabbit anti-Iba1, 1:500, Cat. No CP290A, BioCare medical; mouse monoclonal anti-HA, 1:100, Cat. No 16B12, BioLegend; rat anti-P2RY12, 1:100; Cat. No 84001, BioLegend). After washes in PBS, sections were incubated for 2 hours at room temperature with a secondary antibody diluted in the blocking solution (goat Alexa Fluor 594-conjugated anti-rabbit IgG, 1:1000, Cat. No A11037, LifeTechnologies; donkey CF488A-conjugated anti-mouse IgG, 1:1000, SAB4600035, Sigma; goat Alexa Fluor 633-conjugated anti-rat IgG, 1:1000, Cat. No A-21094, Invitrogen). After further washes in PBS, sections were incubated with Hoechst (1:2000 in blocking solution, Cat. No H3570, Invitrogen) for 10 min at room temperature. Sections were washed several times, finally held with mounting medium (Fluoromount; Cat. No S3025, Dako), coverslipped, and stored at 4°C until use. Acquisitions were done with a confocal laser-scanning microscope (SP8, Leica). For morphological analysis of Iba1-positive microglia, a X63 oil-immersion objective was used and Z-stack interval was 0.30 µm.

### Image analysis

All image analyses were carried out blindly, i.e. without disclosing the experimental group of mice. Confocal image stacks were analyzed using Imaris software (v. 9.9.1). Surface rendering was done for each channel. Total process length, number of branch points, and number of Sholl intersections were determined using the Filament function.

### Brain tissue dissociation and separation for FACS analysis

Cell purification flowchart from brain tissue dissociation to FACS cell sorting is shown in Extended Data Fig. 5c. After intracardiac perfusion with a buffered solution (free HBSS w/o Ca^2+^, Mg^2+^ and phenol red), brain or hypothalamus were collected, cut in small pieces with scissors and subjected to enzymatic and mechanical cell dissociation as previously described ^61^. Small pieces of tissues were incubated for 15 min at 22°C in HBSS (10 ml per brain or 3 ml per hypothalamus) containing 0.05% collagenase D (Roche), 0.1 µg/ml N-α-tosyl-L-lysine chloromethyl ketone hydrochloride (TLCK, Sigma), 0.4% dispase II (Roche), 0.025 U/ml DNaseI (Sigma-Aldrich) and 10 mM HEPES buffer (Invitrogen). Pieces of tissues were dissociated with a glass dounce homogenizer equipped with an A pestle and further incubated for 15 min at 22°C. Homogenized tissues were filtered through a 70 µm cell strainer (Cat. No 93070, SPL Life Sciences). Then, cell suspensions were centrifuged (7 min, 300 g) and washed. Cells from whole brain (not from hypothalamus) were further separated over discontinuous Percoll gradients. Cells from a quarter brain were resuspended in 14 ml of 67.5% isotonic Percoll (Percoll® 67.5%, HBSS w/o Ca^2+^, Mg^2+^ and phenol red, 0.14% glucose, 2% BSA and 3,625 mM HEPES) and transferred to 50 ml polypropylene conical tubes with equal volumes of 22.5% isotonic Percoll (22.5% Percoll®, 1x HBSS, 1.65% BSA) and 10 ml of PBS 1x sequentially layered on top. After centrifugation (800 g, 25 min, 4°C), the 22.5%:0% Percoll interphase containing myelin was removed and the 67.5% :22.5% Percoll interphase containing macrophages and microglial cells was collected. Cells were centrifuged (500 g, 10 min), washed in PBS, and then immunolabeled for flow cytometry analysis or sorting.

### FACS cell sorting

Dissociated and purified cells from the brain without cerebellum were incubated in 10% mouse serum (diluted in 1x PBS, 1% BSA) to block Fc receptors, and immunostained with following fluorophore-conjugated antibodies: rat anti-CD11b (PE-Cy™7 conjugated, 1:80; Cat. No 552850, BD Pharmingen) and rat anti-CD45 (BV421 conjugated, 1:100; Cat. No 563890, BD Horizon), plus a fixable viability stain (Cat. No 564405, BD Biosciences) for live/dead cell discrimination. Microglial (CD11b^+^/CD45^low^) and not microglial (CD11b^-^) cells were sorted on a BD FACSAria^TM^ II cell sorter flow cytometer following the gating strategy shown in Extended Data Fig. 5d. Peripheral blood was collected by intracardiac puncture on anesthetized mice in EDTA coated tubes. Samples were treated with a lysis buffer for 15 min at 22°C for lysing red blood cells (BD FACS™ Lysing solution; Cat. No 349202, BD Biosciences). Remaining white blood cells were blocked with 10% mouse serum (diluted in 1x PBS, 1% BSA), and immunostained with rat anti-CD11b (PE-Cy™7 conjugated, 1:80; Cat. No 552850, BD Pharmingen) and rat anti-Ly6G (FITC-conjugated, 1:10; Cat. No 130102296, MACS Miltenyi). Monocytes were identified based on Ly6G^-^/CD11b^+^/SSC^low^ expression ^62^, and were sorted on a BD FACSAria^TM^ II cell sorter flow cytometer using the gating strategy shown in Extended Data Fig. 5e.

### Analysis of MBH cell populations by flow cytometry

Dissociated and purified cells from MBH were analyzed to quantify the percentage of microglia, macrophages, granulocytes, and lymphoid cells in the hypothalamus. Flow cytometry assessed the cellular morphological parameters of size (Forward Scatter, FSC) and granularity (Side Scatter, SSC), and the expression level of surface markers using antibodies. Cells were incubated with 10% mouse serum (in 1x PBS, 1% BSA) for blockade of Fc receptors and were immunostained with following antibodies and reagents: rat anti-CD11b (PE-Cy™7 conjugated, 1:80; Cat. No 552850, BD Pharmingen), rat anti-Ly6G (FITC-conjugated, 1:10; Cat. No 130102296, MACS Miltenyi), Ly6C (rat anti-Ly6C vioGreen conjugated, 1:10; Cat. No 130102207, MACS Miltenyi), rat anti-CD45 (BV421 conjugated, 1:100; Cat. No 563890, BD Horizon), rat anti-CD206 (BV711 conjugated, 1:80; Cat. No 1308635, Sony Biotechnology), rat anti-P2RY12 (Biotin conjugated, 1:50; Cat. No 848007, Biolegend), mouse streptavidin (APC-conjugated, 1:400; Cat. No 2626035, Sony Biotechnology). VersaComp Antibody Capture Kit (Cat. No B22804, Beckman Coulter) was used to set compensation. Cells were acquired with a flow cytometer (BD LSRFortessa^TM^) and analyzed with FlowJo^TM^ (BD Biosciences, version V10.0.7r2) using the gating strategy shown in Extended Data Fig. 1j, which identified microglia (CD11b^+^-CD45^low^-CD206^-^-P2RY12^high^), macrophages (CD11b^+^-CD45^high^-CD206^+^-P2RY12^int^), granulocytes (CD11b^+^-CD45^+^-Ly6G^+^-Ly6C^+^) and lymphoid cells (CD11b^-^/CD45^+^) (Extended Data Fig. 1k).

### Primary culture of microglia

Brains from five to eight male mice (2-3 days old) from Cx3cr1^CreER/+^-*Asc*^fl/fl^, *Cx3cr1*^CreER/+^-*Asc*^wt/wt^, *Asc*^KO^ or C57BL6/J mice were dissected. Both hemispheres were separated and the surrounding meninges were removed in PBS, under a binocular magnifier. The cleaned brains were dissociated into a single-cell suspension by tissue trituration with a 18-gauge then a 21-gauge syringe. Cells were pelleted at 300 g for 10 min at 4°C and the pellet resuspended in fresh complete culture medium (DMEM 4.5 g/l of glucose, #41965-039, Gibco; 10% heat-inactivated Fetal Bovine Serum, #A5256701, Gibco; 100u/mL Penicillin-100µg/mL Streptamycin, #15140-122, Gibco). Cells obtained from each individual brain were plated on poly-L-lysine-coated 75cm^2^ flasks in fresh complete culture medium at 37°C and 5% CO_2_ to generate mixed glial cultures. 5 and 10 days later, the complete culture medium was renewed and on day 14-16, isolation of microglia was performed using a magnetic cell sorting system to positively select microglia that are CD11b^+^. Confluent mixed cells were detached from the flasks, magnetically labelled with anti-CD11b MicroBeads (#130-093-636, Miltenyi Biotec) and loaded on a LS column (#130-042-401, Miltenyi Biotec) placed into an OctoMACS™ Separator (#130-042-108, Miltenyi Biotec) according to the manufacturer’s instructions. The columns were washed with 3 × 3 ml PBS 1X / 0.5% BSA (PBS-BSA). The columns were removed from the magnetic field, and microglia were flushed out using 5 ml PBS-BSA. To obtain a higher purity, the eluate containing microglia was applied onto a MS column (#130-042-201, Miltenyi Biotec) placed into the OctoMACS^TM^ Separator (#130-042-108, Miltenyi Biotec). The columns were washed with 3 × 0.5 ml PBS-BSA and microglia were flushed out using 1 ml PBS-BSA. Isolated microglia were cultured in mixed cell culture-conditioned medium ASC protein-free or not, obtained from mixed cell culture prepared from brain of Cx3cr1^CreER/+^-*Asc*^wt/wt^ or *Asc*^KO^ mice. Cells were seeded at 150,000 cells/cm^2^ in poly-L-lysine-coated plates for anti-ASC immunolabelling or at 30,000 cells/cm^2^ for *Asc* genotyping and kept at 37°C and 5% CO_2_. The mixed cell culture-conditioned medium corresponds to the culture medium recovered from the mixed glial cell cultures at day 14-16. Twenty-two hours after magnetic cell sorting, the isolated microglia (from *Cx3cr1*^CreER/+^-*Asc*^fl/fl^ mice or control *Cx3cr1*^CreER/+^-*Asc*^wt/wt^ mice) were incubated with 2.5 µM 4-hydroxy-tamoxifen for 72 h to induce *Asc* deletion. Medium was changed, cells were kept for additional 4 days then *Asc* genotyping and ASC immunolabelling were performed.

ASC speck formation was induced by LPS priming (Lipopolysaccharides from Escherichia coli O127:B8) (1µg/ml, #L3129, Merck^TM^) for 3 h followed by treatment with 10 µM Nigericin (#N1945, Merck^TM^) for 30 min. Cells were rinsed in PBS and fixed with 4% paraformaldehyde for 20 min for subsequent immunolabeling.

Fixed cells were washed twice in PBS and then permeabilized in FACS permeabilizing solution (#554714, BD Biosciences) for 10 min at room temperature followed by 3 rinses with PBS1X / Saponin 0.05% (PBSS). Cells were incubated for 20 min at room temperature in a blocking solution (5% mouse serum in PBSS). Cells were then incubated for 20 min at room temperature with primary antibodies (chicken anti-Iba1, 1:1000; Cat. No 234 009, Synaptic systems; rabbit anti-ASC, 1:800; Cat. No D2W8U, Cell Signaling) and for 20 minutes at room temperature with secondary antibodies (AlexaFluor^TM^ 555 goat anti-chicken IgY, 1:1,000, #A21437, Invitrogen; AlexaFluor^TM^ 488 goat anti-rabbit IgG, #A11034, Life Technologies). All antibody incubations were performed using PBS1X, 0.05% saponin, 2% mouse serum. Wash steps were performed using PBSS. Cells were washed and then incubated in Hoechst 33342 (2 µg/ml, 5 min; #3570, Invitrogen,) at room temperature before final washing and mounting (#S3025, DAKO) before imaging using an Axio Imager microscope (Zeiss) equipped with an AxioCam camera (Zeiss).

### Primary culture of peritoneal macrophages

Mice were anesthetized by ip injection of a ketamine – xylazine mix (100 mg/kg – 10 mg/kg). Peritoneal macrophages were collected in complete culture medium (RPMI, 1% penicillin-streptomycin, 10% SVF, 1 mM sodium pyruvate), and plated overnight at 37°C, 5% CO_2_. After extensive washing in PBS to eliminate floating and dead cells, peritoneal macrophages were used.

### Primary culture of bone-marrow-derived-macrophages

Mice were anesthetized by ip injection of a ketamine – xylazine mix (100 mg/kg – 10 mg/kg) and euthanized by decapitation. Femurs and tibias were harvested, and bone marrow cells were flushed out using PBS. Cell suspensions were filtered through a 70 µm cell strainer, and centrifuged (10 min, 350 g, 4°C). Supernatants were discarded, and cells were collected in a complete culture medium (RPMI, 1% penicillin-streptomycin, 10% SVF). Cells were seeded at a density of 1×10^6^ cells in 12-well plates, and plated at 37°C, 5% CO_2_. Differentiation into macrophages was induced by adding recombinant M-CSF (130-096-492, Miltenyi, 0.02%) for 7 days until complete differentiation. Complete culture medium was renewed on day 5, when the cells became adherent. On day 7, differentiated macrophages were gently washed with 3 × 2 ml PBS to eliminate floating and dead cells. For mRNA extraction, cells were lysed with RLT buffer from the RNeasy Micro Kit (Cat. No #74004, Qiagen). Samples were stored at -80°C until use.

### Genotyping

Genotyping was performed on FACS-sorted cells and cultured peritoneal macrophages. Genomic DNA was extracted using the Quick extract DNA extraction solution (Cat. No #QE0905T, Epicentre) according to manufacturer’s protocol. Samples were stored at -20°C until use. The triple primer-PCR analysis generated the *Asc*^deleted^ (∼450 bp), *Asc*^floxed^ (∼520 bp), *Cx3cr1*^Cre^ (∼300 bp) and *Cx3cr1*^WT^ (∼750 bp) amplicons. PCRs were carried out in 20 µl reaction volume containing 1 µl of DNA, 10 µl of KAPA mouse genotyping kit master mix (Cat No. KK5621, Cliniscience), nuclease-free water and appropriate probes. PCR targeting the *Asc* gene were performed using the following primers: ASC for (5’GGCTTAGTGTTCTAGCTGTT3’; 1 µl), ASCrevWT (5’ACCAGGAAGTCAGCTGCTTA3’; 1 µl), and ASCspe (5’TTCACACAATACGATCTTCA3’; 2 µl). Reactions were performed on a SimplyAmp Thermal Cycler (Applied Biosystems) with the following PCR cycling conditions: 95°C for 2 min followed by 35 to 40 cycles, including 94°C for 7 sec, 56.5°C for 20 sec, and 72°C for 20 sec, and a final step 72°C for 3 min. PCR targeting the Cx3cr1 gene were performed using the following primers: CX3for (5’CCTCTAAGACTCACGTGGACCTG3’; 2 µl), CX3revWT (5’GACTTCCGAGTTGCGGAGCAC3’; 2 µl), and CX3spe (5’GCCGCCCACGACCGGCAAAC3’; 1 µl). Reactions were performed on a SimplyAmp Thermal Cycler (Applied Biosystems) with the following PCR cycling conditions: 95°C for 3 min followed by 35 to 40 cycles, including 94°C for 15 sec, 60°C for 15 sec, and 72°C for 20 sec, and a final step 72°C for 5 min.

### RNA isolation and RT-qPCR from FACS-sorted cells and cultured cells

mRNA analysis of FACS-sorted cells and peritoneal macrophages and BMDM were performed on total RNA which were extracted using the RNeasy Micro kit (Qiagen) according to the manufacturer’s instructions. mRNA analyses of tissue were performed on total RNA extracted from brain, liver, or spleen tissue after grinding the tissue using the Tissue Lyser II (Qiagen). Total RNA was extracted using the NucleoSpin RNA Plus kit (#740984, Macherey-Nagel) according to the supplier’s instructions. The concentration and the integrity of extracts were assessed by spectrophotometry (Nanodrop 2000; ThermoFisher Scientific). The mRNA samples were stored at -80°C until use. Reverse transcription was performed on 200 ng of mRNA using SuperScript IV VILO Master Mix (Cat No. 11766050, ThermoFisher). Reactions were performed on a SimplyAmp Thermal Cycler (Applied Biosystems) with the following cycling conditions: 25°C for 10 min, 50°C for 10 min, and 85°C for 5 min. cDNA samples were stored at -20°C until use. qPCRs were carried out in a 10 µl reaction volume containing 2 µl of 1:10 diluted cDNA, 2 µl of nuclease-free water, 5 µl of Taqman fast advanced Master Mix (Applied Biosystems), and 1 µl of appropriate Taqman probes (ThermoFisher Scientific) for Polr2a (Mm00839502) and *Asc* (Mm00445747). Real-time PCRs were run on a StepOnePlus thermocycler (Applied Biosystems) with the following cycling conditions: 95°C for 2 min, 40 cycles including 95°C for 1 sec, and 60°C for 20 sec. The comparative cycle threshold method was used for relative quantification of target genes, using Polr2a as a reference gene. The Ctrl genotype was used as controls in the analysis.

### Polyribosome immunoprecipitation

Mice were deeply anesthetized by intraperitoneal injection of a ketamine – xylazine mix (100 mg/kg – 10 mg/kg), and transcardiacally perfused with PBS (15 ml). Brains were collected, MBH (Medio Basal Hypothalamus) were dissected and quick frozen at -80°C in liquid nitrogen, and stored at -80°C until use. 5 biological replicates, each composed of a pool of 6 mice, were used for RNAseq. HA-tagged ribosome immunoprecipitation was performed as described previously^32^ with slight modifications. The hypothalamus from mice was homogenized by douncing in 1 ml polysome buffer (50 mM Tris, pH 7.4, 100 mM KCl, 12 mM MgCl₂, and 1 percent NP-40 supplemented with 1 mM DTT, 1 mg/ml heparin, 100 μg/ml cycloheximide, 200 U/ml RNaseOUT, and protease inhibitor mixture). Samples were then centrifuged at 10,000 × g for 10 minutes, and the supernatant was collected in a new Eppendorf tube. A 50 μl aliquot of each supernatant was transferred to a separate tube to serve as the input fraction for validation. Anti-HA antibody (4 μl/sample; BioLegend cat. 16B12) was added to the remaining supernatant and incubated overnight at 4 °C with gentle constant rotation. The following day, samples were added to 200 μl of protein G magnetic beads (Invitrogen, #100.04D), which had been pre-washed twice with 200 μl of homogenization buffer (50 mM Tris, pH 7.4, 100 mM KCl, 12 mM MgCl₂, and 1 percent NP-40), and incubated overnight at 4 °C with gentle constant rotation. On the third day, magnetic beads were washed twice on a magnetic rack for 10 minutes each with 800 μl of high-salt buffer (50 mM Tris, pH 7.4, 300 mM KCl, 12 mM MgCl₂, 1 percent NP-40, 1 mM DTT, and 100 μg/ml cycloheximide). After washing, 350 μl of Qiagen RLT buffer were added to the pellets and to the input samples. Samples were immediately frozen on dry ice and stored at - 80 °C until use. Total RNA was extracted and purified using the RNeasy Plus Micro Kit (Qiagen, #74004) according to the manufacturer’s instructions. RNA was eluted in RNase/DNase-free water and stored at -80 °C until use. RNA yields from the HA-tagged ribosome immunoprecipitates were quantified using the Agilent 2100 Bioanalyzer (Agilent Technologies). RNA integrity was assessed for all samples using the Agilent 2100 Bioanalyzer, and all RNA integrity numbers (RIN) were above 7.0.

### RNA sequencing

Library preparation and RNA sequencing were carried out by the MGX core facility (Biocampus, Montpellier, France). In brief, cDNA libraries were generated from 4 ng total RNA using the SMART-Seq total RNA Pico Input with UMIs (Takara bio), following the manufacturer’s recommendations. Sequencing was performed on an Illumina NextSeq 500 system using an SP flow cell in 50 bp paired-end mode. Across samples, sequencing depth ranged from 57 to 96 million reads, and all libraries successfully passed the facility’s quality-control metrics.

### Bioinformatics

After RNA sequencing, clean raw counts were aligned against Mus musculus reference genome version GRCm39 with STAR and quantified with Salmon through a nf-core/RNAseq pipeline v3.14.0^63^. Raw counts were normalized using R software (R core Team, 2025) and the edgeR package^64^. Gene expression levels were analyzed using gene-wise linear regression models. For each gene, expression values were modeled independently across samples using ordinary least squares regression. Four alternative models were considered in order to test different hypotheses regarding the relationship between gene expression, lipid availability, and dietary group. First, a model without any explanatory variable (y_i_=β_0_+ε_i_). Second, a linear model including lipid quantity (y_i_=β_0_+L_i_+ε_i_). Third, a group-based model (y_i_=β_0_+ β_Gi_+ε_i_). Finally, a quadratic model was used to capture potential non-linear relationships between gene expression and lipid quantity (y =β + β L + β L ^2^+ε). For each gene, model comparison was performed using the Bayesian Information Criterion (BIC). The model with the lowest BIC value was selected as the best-supported model. PCA was calculated using FactoMineR package ^65^. Hierarchical clustering was applied to the genes associated to the dimension 1 of the PCA (cos^2^>0.7) using the 1-Pearson correlation coefficient as distance and Ward’s criterion for agglomeration. The clustering results are presented using the pheatmap package (https://CRAN.R-project.org/package=pheatmap) as a heatmap of normalized log_2_ counts per million, centered and scaled. GSEA was performed using the fgsea package (https://doi.org/10.1101/060012), with the loadings of the genes on the first dimension of the PCA as the score. Gene Ontology categories networks were plotted using ggraph and igraph packages (https://CRAN.R-project.org/package=ggraph; https://doi.org/10.5281/zenodo.17470081). A link was drawn between two categories if a smaller category shares 90% homology in its leading edges genes composition with a larger category. Volcano plots and dot plots of gene expression data were realized with the tidyverse package ^66^.

### Statistical analysis

Statistical analyses were performed using GraphPad Prism software (v. 10.2.2, GraphPad Software Inc.). No statistical method was used to predefine sample size. Results on graphs are shown as means +/- standard errors of the mean (SEM). No correction was applied on data for analyses. Equality of variances and normality of distribution were checked prior analysis. Comparisons between 2 groups were carried out using two-tailed Student test or Mann-Whitney test. Multiple comparisons of groups were carried out using one-way analysis of variance (ANOVA) followed by post-hoc Tukey test or using nonparametric Kruskal-Wallis test followed by post-hoc Dunn test, as appropriate and indicated in legends. To analysis effect of two independent factors, two-way ANOVA was used, followed by the post-hoc Sidak’s test.

## Extended data

**Suppl Figure 1.**
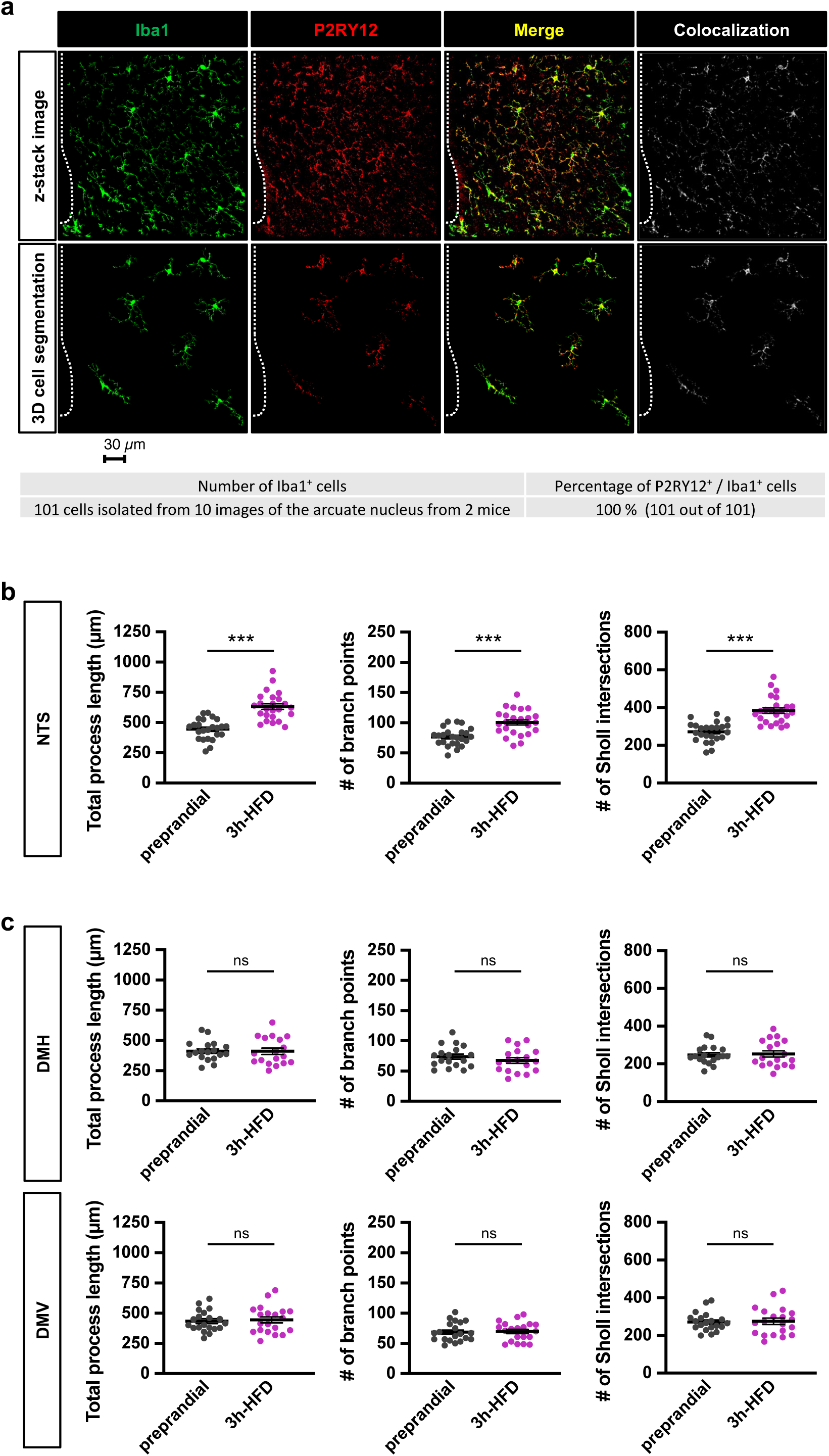
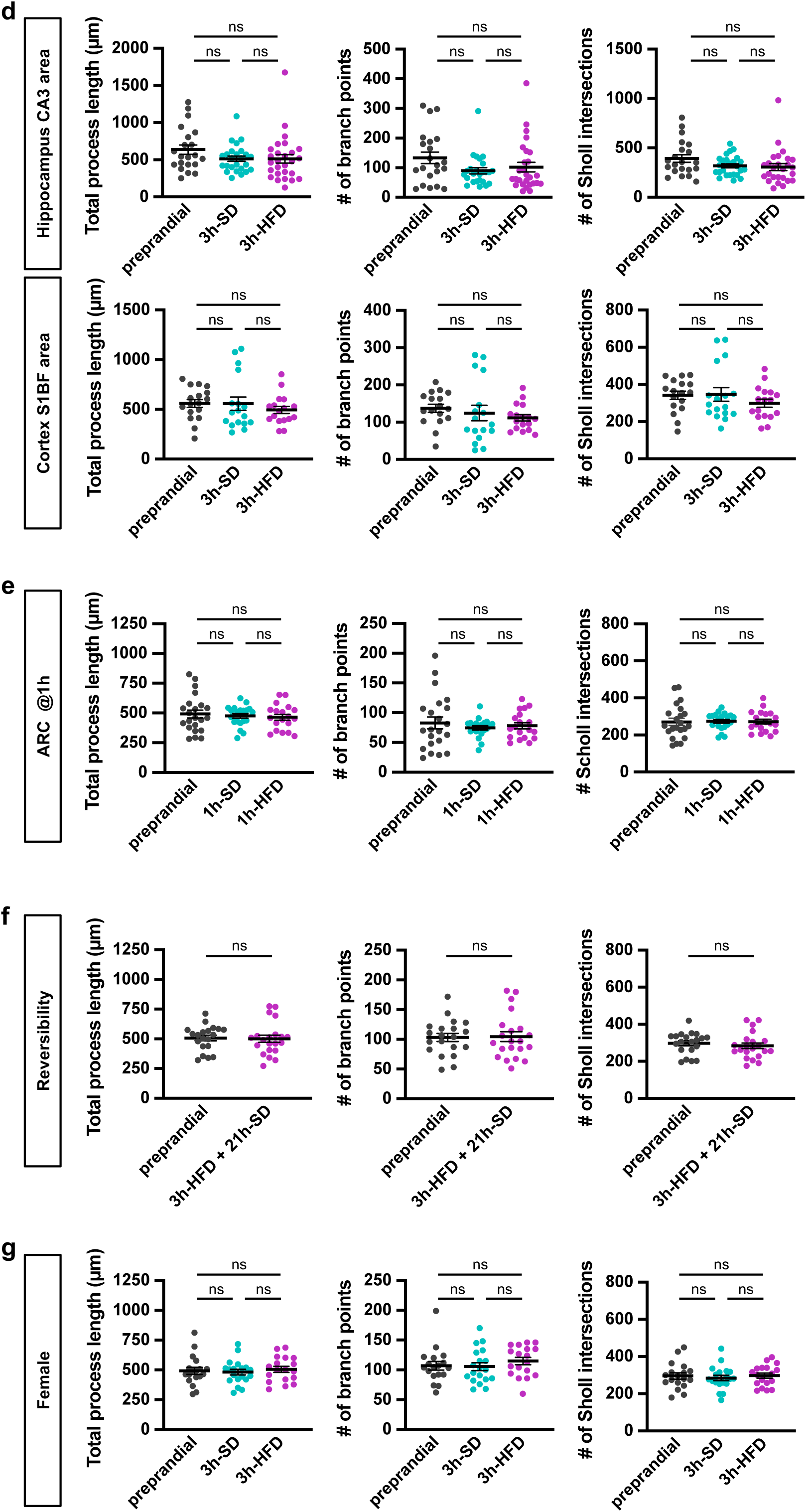
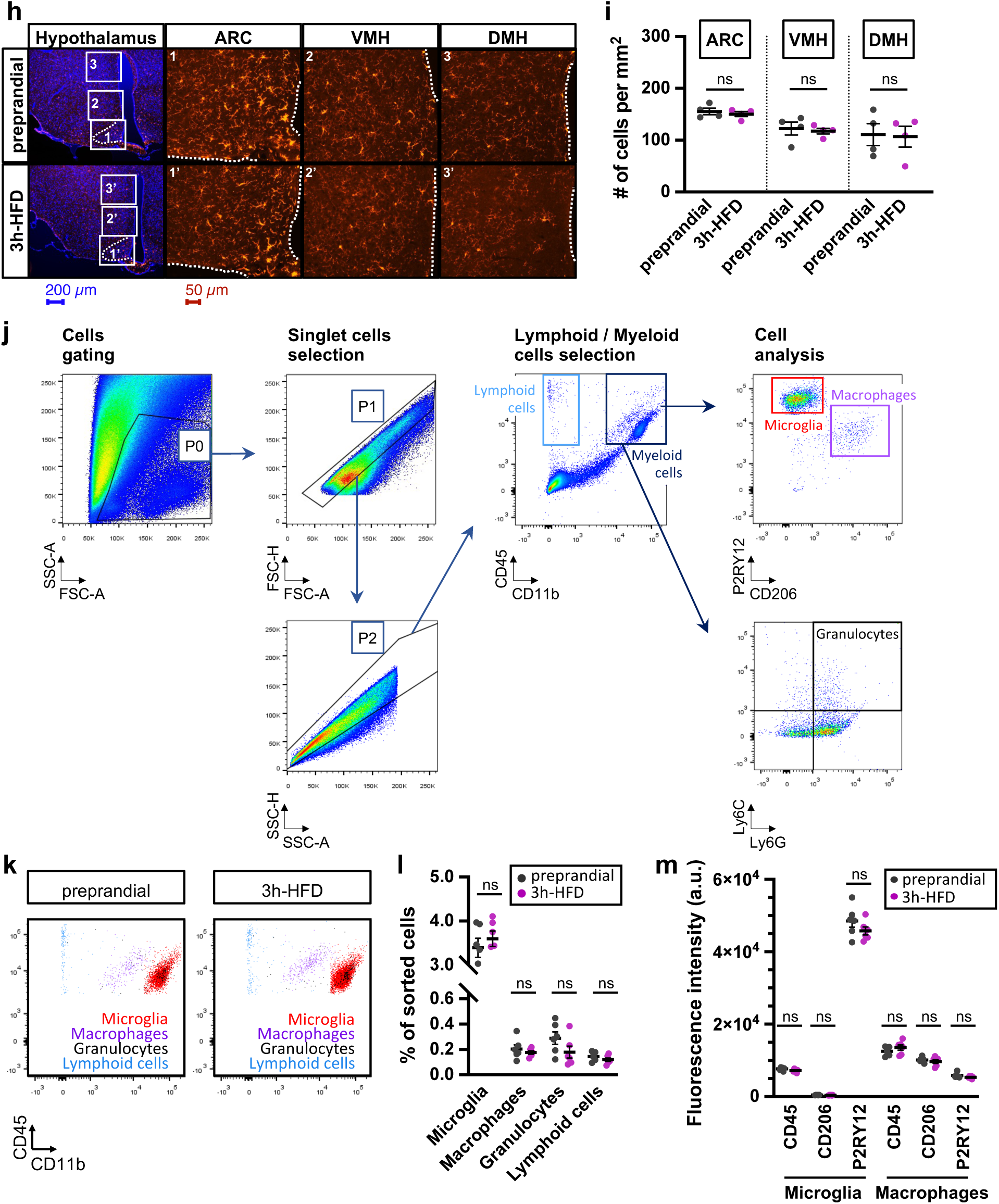
Regioselectivity, reversibility and sex-specificity of the HFD-induced postprandial microglial reactivity. **(a)** Microglia of the mediobasal hypothalamus from fixed brain slices of 8-week-old C57BL/6J male mice were stained using anti-Iba1 (in green) and anti-P2RY12 (in red) antibodies. Images were taken with a confocal microscope and analysed using Imaris software. In **(b-g)**, morphology of brain microglia was examined in 8/10-week-old C57BL/6J mice in the preprandial state or fed with either standard diet (SD) or high fat diet (HFD). Iba1^+^ microglia were immunodetected in fixed brain slices using anti-Iba1 antibodies and imaged by confocal microscopy. Iba1^+^ cells were reconstructed in 3-dimensions using Imaris software. Morphometric characteristics of Iba1^+^ cells were computed with Imaris software. In **(b-d)**, brain slices were obtained from male mice in the preprandial state or fed with either standard diet (SD) or high fat diet (HFD) for 3h. Analyses were done in the nucleus of the solitary tract (NTS) in the brainstem **(b)**, in the dorsomedial hypothalamus (DMH) and the dorsal motor nucleus of the vagus (DMV) in the brainstem **(c)**, in the CA3 area of the hippocampus and in the S1BF cortical area **(d)**. In **(e)**, brain slices were obtained from male mice in the preprandial state or fed with either standard diet (SD) or high fat diet (HFD) for 1h only. Analyses were done in the arcuate nucleus of the hypothalamus (ARC). In **(f)**, brain slices were obtained from male mice in the preprandial state or fed with high fat diet (HFD) for 3h and then with standard diet (SD) for 21h. Analyses were done in the arcuate nucleus of the hypothalamus (ARC). In **(g)**, brain slices were obtained from female mice in the preprandial state or fed with either standard diet (SD) or high fat diet (HFD) for 3h. **(h)** Microscopic images of Iba1^+^ cells in the hypothalamus from fixed brain slices of 8/10-week-old C57BL/6J male mice in the preprandial state or fed with a high fat diet (HFD) for 3h. Images show red fluorescent Iba1^+^ cells in the arcuate nucleus (ARC), ventromedian hypothalamus (VMH) and dorsomedian hypothalamus (DMH). **(i)** Quantification of microglia number in the hypothalamus of mice according to the prandial state (preprandial, n= 4 mice; 3h-HFD, n= 4 mice). Cell counting was performed using microscopy. Iba1^+^ cells in fixed brain slices were counted on microscopic images using Image J software. Counting was performed in the arcuate nucleus (ARC), the ventromedial hypothalamus (VMH), and the dorsomedial hypothalamus (DMH). **(j)** Gating strategy of FACS-sorted microglia, macrophages, granulocytes, and lymphoid cells from the mediobasal hypothalamus of 9/10-week-old C57BL/6J male mice. The population of interest cleared of cell debris (P0) was selected on the forward scatter-area (FSC-A) versus side scatter-area (SSC-A) density plot. Single cells were gated (population P2) using the forward scatter-height (FSC-H) versus forward scatter-area (FSC-A) and the side scatter-height (FSC-H) versus side scatter-area (FSC-A) density plots. Further cell identification was performed using CD11b, CD45, P2RY12, CD206, Ly6G and Ly6C markers. Myeloid cells including microglia, macrophages, and granulocytes were identified as (CD11b^+^/CD45^+^), lymphoid cells were identified as (CD11b^-^/CD45^+^). More specifically, microglia were identified as (CD11b^+^/CD45^low^/P2RY12^high^/CD206^-^), macrophages as (CD11b^+^/CD45^high^/P2RY12^int^/CD206^+^), and granulocytes as (CD11b^+^/CD45^+^/Ly6G^+^/Ly6C^+^). **(k)** Representative flow cytometry profiles from dissociated hypothalamic cell suspensions prepared from mice in the preprandial state or fed with HFD for 3 hours. **(l)** Percentage of microglia, macrophages, granulocytes, and lymphoid cells in the mediobasal hypothalamus of mice according to the prandial state (preprandial, n= 5-6 mice; 3h-HFD, n= 5-6 mice). Cell counting was performed using flow cytometry. Cells were analysed from dissociated hypothalamic cell suspensions. Percentages were calculated for each cell population relative to the total P2 gated hypothalamic cell population (see Extended Data Fig. 1j) (preprandial, n=6 mice; 3h-HFD, n=6 mice. **(m)** Quantification of CD45, CD206 and P2RY12 mean fluorescence intensity (a.u.) in microglia and macrophages in the hypothalamus of mice according to the prandial state. **Statistics.** Data are mean ± standard error of the mean (SEM). Asterisks indicate statistical differences between groups: ***p<0.001; ns, non-significant. Data were analysed using two-tailed unpaired Student’s test (in b, c, f), Kruskal-Wallis and Dunn’s tests (in d, e, g), and two-way ANOVA and Tukey’s tests (in i, l, m). 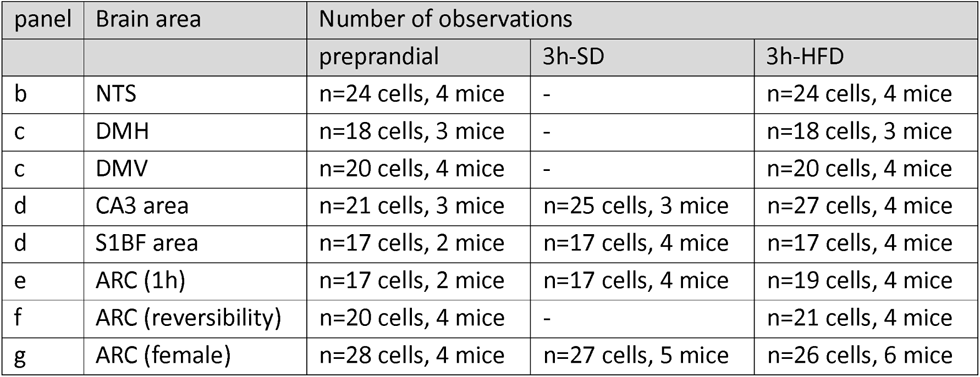

**Suppl Figure 2.**
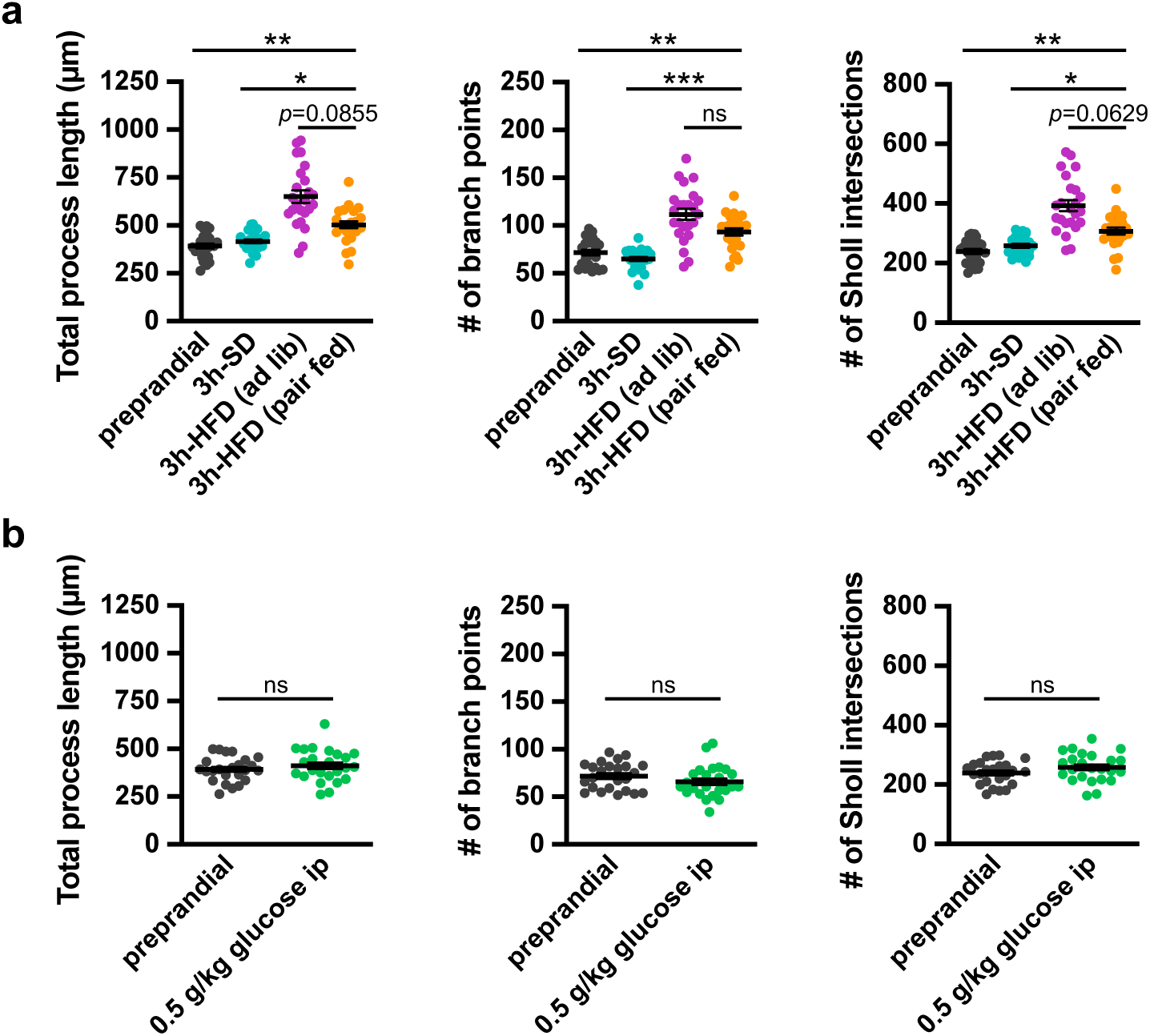
Characteristics of the postprandial microglial reactivity. In **(a, b)**, morphology of brain microglia was examined in 8/10-week-old male C57BL/6J mice according to the prandial state. Experimental groups of mice were: preprandial mice (no food); postprandial mice, in which mice were fed with SD or HFD for 3 hours (3h-SD and 3h-HFD); postprandial pair-fed mice, in which the amount of HFD food provided to the mice was matched to that consumed by the 3h-SD group (i.e. 0.8 g of HFD); and postprandial mice, in which the food was replaced by a bolus of glucose that mimics the postprandial hyperglycemia (ip injection of 0.5 g/kg glucose). The five experimental groups of mice have been experienced all together at the same time. After sacrifice, Iba1^+^ microglia were immunodetected in fixed brain slices using anti-Iba1 antibodies and imaged by confocal microscopy. Iba1^+^ cells were reconstructed in 3-dimensions using Imaris software. Morphometric characteristics of Iba1^+^ cells were computed with Imaris software. Suppl Figure 2a and b includes data from main figure 1e for comparison and statistical purposes. **(a)** Morphometric characteristics of arcuate Iba1^+^ cells according to the prandial state (preprandial, n=24 cells from 4 mice; 3h-SD, n=24 cells from 4 mice; 3h-HFD ad libitum, n=24 cells from 4 mice; 3h-HFD pair fed, n=24 cells from 4 mice). **(b)** Morphometric characteristics of arcuate Iba1^+^ cells in glucose-injected mice compared to preprandial mice (preprandial, n=24 cells from 4 mice; glucose, n=24 cells from 4 mice). **Statistics.** Data are mean ± standard error of the mean (SEM). Asterisks indicate statistical differences between groups: *p<0.05; ***p<0.01; ***p<0.001; ns, non-significant. Data were analysed using one-way ANOVA and Dunn’s (in a), two-tailed unpaired Student’s test (in b).

**Suppl. Figure 3.**
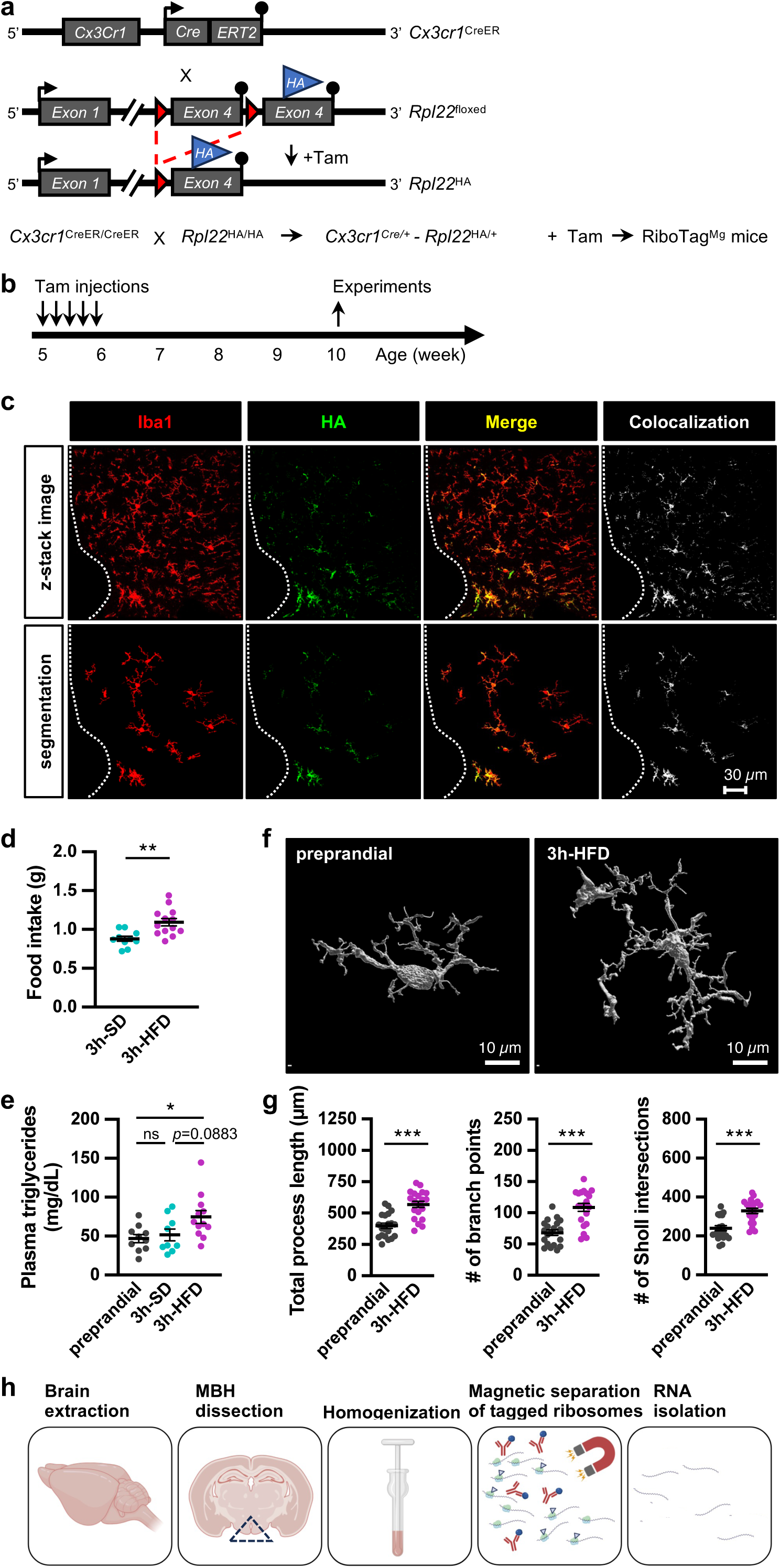
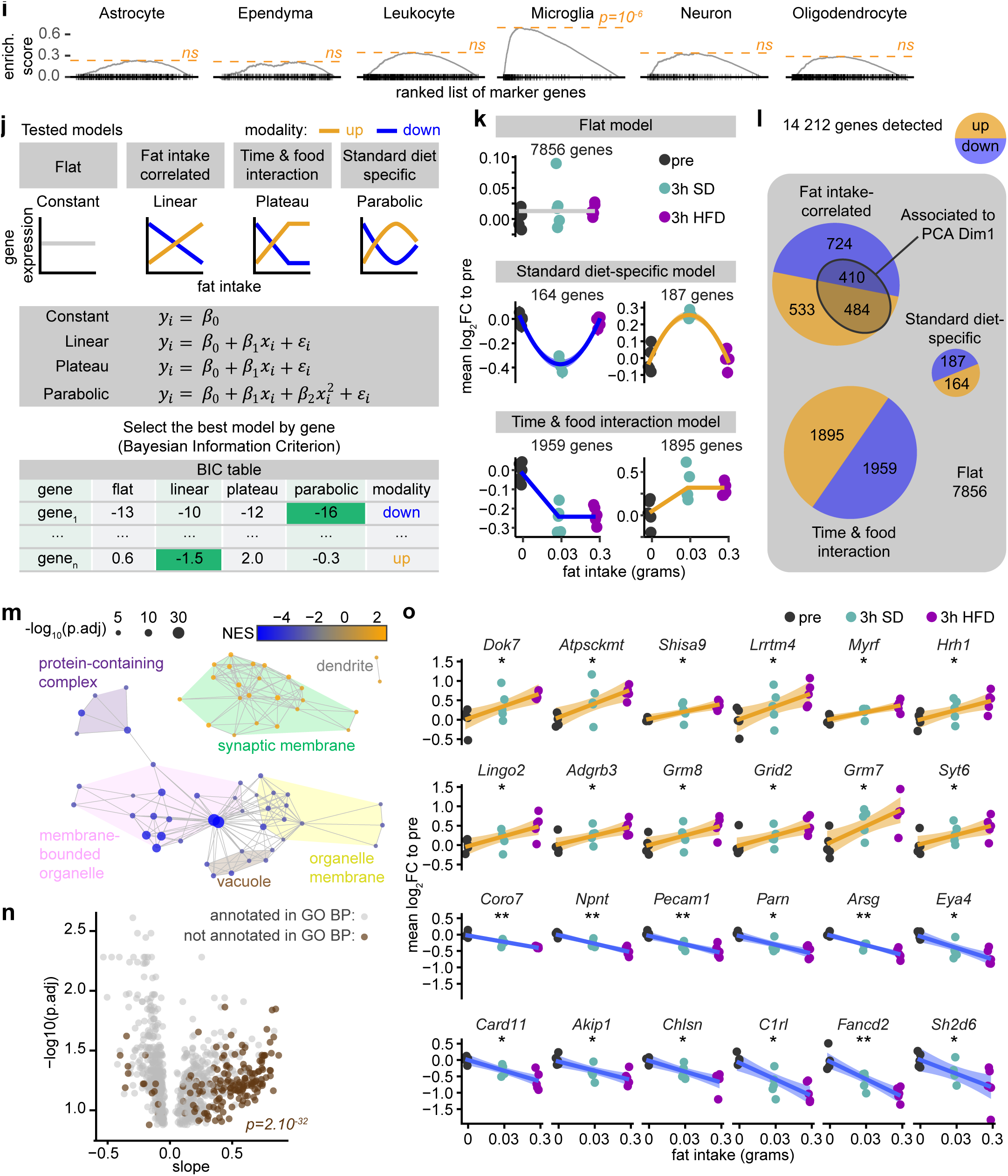
Generation and validation of the microglia-specific RiboTag mouse model (RiboTag^Mg^ mice), and high-throughput RNA sequencing analysis of the microglial translatome according to the prandial state. **(a)** Schematic showing the strategy used to express the Rpl22^HA^ gene in Cx3cr1^+^ cells. Conditional gene targeting was based on the tamoxifen-inducible Cre-lox system driven by the Cx3cr1 promoter. Cx3cr1^CreER^ mice were crossed with mice harboring a floxed Rpl22^HA^ gene. After tamoxifen treatment, a specific expression of the Rpl22^HA^ gene occurred in Cx3cr1^+^ cells. **(b)** Schematic showing protocol of tamoxifen treatment used to induce Rpl22^HA^ expression in microglia. 5-week-old Cx3cr1^CreER/+^:Rpl22^HA/+^ mice were injected with tamoxifen (75 mg/kg) for 5 consecutive days. Further experiments were carried out on 10-week-old tamoxifen-treated Cx3cr1^CreER^:Rpl22^HA^ mice (i.e. RiboTag^Mg^ mice). **(c)** Expression of Hemagglutinin (HA) epitope-tagged RPL22 ribosomal protein in Iba1^+^ Cx3cr1-expressing microglial cells within the hypothalamus in fixed brain slices from RiboTag^Mg^ mice. Immunodetection was done using anti-Iba1 antibody (red), and anti-HA antibody (green). **(d)** Food intake of RiboTag^Mg^ mice fed for 3 hours with either standard (3h-SD) or high fat diet (3h-HFD) (3h-SD: n= 10 mice; 3h-HFD: n= 13 mice). **(e)** Plasma triglycerides level after a 3-hour-period of feeding with either SD or HFD (preprandial, n=10 mice; 3h-SD: n= 9 mice; 3h-HFD: n= 12 mice). **(f)** Representative 3D reconstructed images of microglia in the arcuate nucleus of the hypothalamus of mice in the preprandial state or in the postprandial state, upon high fat diet for 3 hours (3h-HFD). Microglia were immunodetected in fixed brain slices using anti-Iba1 antibodies and imaged by confocal microscopy. Iba1^+^ cells were reconstructed in 3-dimensions using Imaris software. **(g)** Morphometric characteristics of arcuate Iba1^+^ cells according to the prandial state (preprandial, n=20 cells from 4 mice; 3h-HFD, n=20 cells from 4 mice). Automatic quantification of cell morphology was done using Imaris software. **(h)** Scheme of the RiboTag methodology including collection of brain from RiboTag mice, dissection of the mediobasal hypothalamus (MBH), tissue homogenization, magnetic isolation of translating polyribosomes using anti-HA antibody coupled with magnetic beads, and RNA extraction. **(i)** GSEA enrichment result of RiboTag data using single-cell data from the brain as database (https://bis.zju.edu.cn). **(j)** Analysis framework. The genes are associated to some pre-determined models using a Bayesian Inclusion Criteria (BIC). The best model is identified as the one with the lower BIC. **(k)** Representation RiboTag data from microglia. Data is mean log2 fold-change to the pre-prandial group of RiboTag data across all identified genes for each individual. SEM is too small to be visible. The blue line represents a negative linear correlation to the log_10_ fat intake, whereas the orange line represents a positive correlation. **(l)** Pie charts of the repartition of RiboTag data throughout the pre-determined models. The areas in orange represents genes that are positively correlated to a model, whereas a blue area represents genes that are negatively associated. **(m)** Representation of RiboTag data from microglia. Data is the log2 fold-change to the pre-prandial group of RiboTag data for each individual. The blue line represents a negative linear correlation to the log_10_ fat intake, whereas the orange line represents a positive correlation. The p value associated to the correlation is indicated : * p <0.05 ; ** p <0.01. **(n)** Volcano plot of genes from the fat intake-correlated model, using the slope and its associated -log10 adjusted p value. Genes in brown do not have a clear annotation in Gene Ontology database. An enrichment of these genes using hypergeometric test is provided (p=2.10^-32^). **(o)** Network of GSEA results of Gene Ontology Cellular Component. A link was drawn between two categories if a smaller category shares 90% homology in its leading edges genes composition with a larger category. The size represents the -log_10_ adjusted p value, and the color the normalized enrichment score. The most central category is used to identify each area of the network. **Statistics.** Data are mean ± standard error of the mean (SEM). Asterisks indicate statistical differences between groups: ***p<0.05; ***p<0.01; ***p<0.001; ns, non-significant. Data were analysed using two-tailed unpaired Student’s test (in d, g), one-way ANOVA and Tukey’s tests (in e).

**Suppl. Figure 4.**
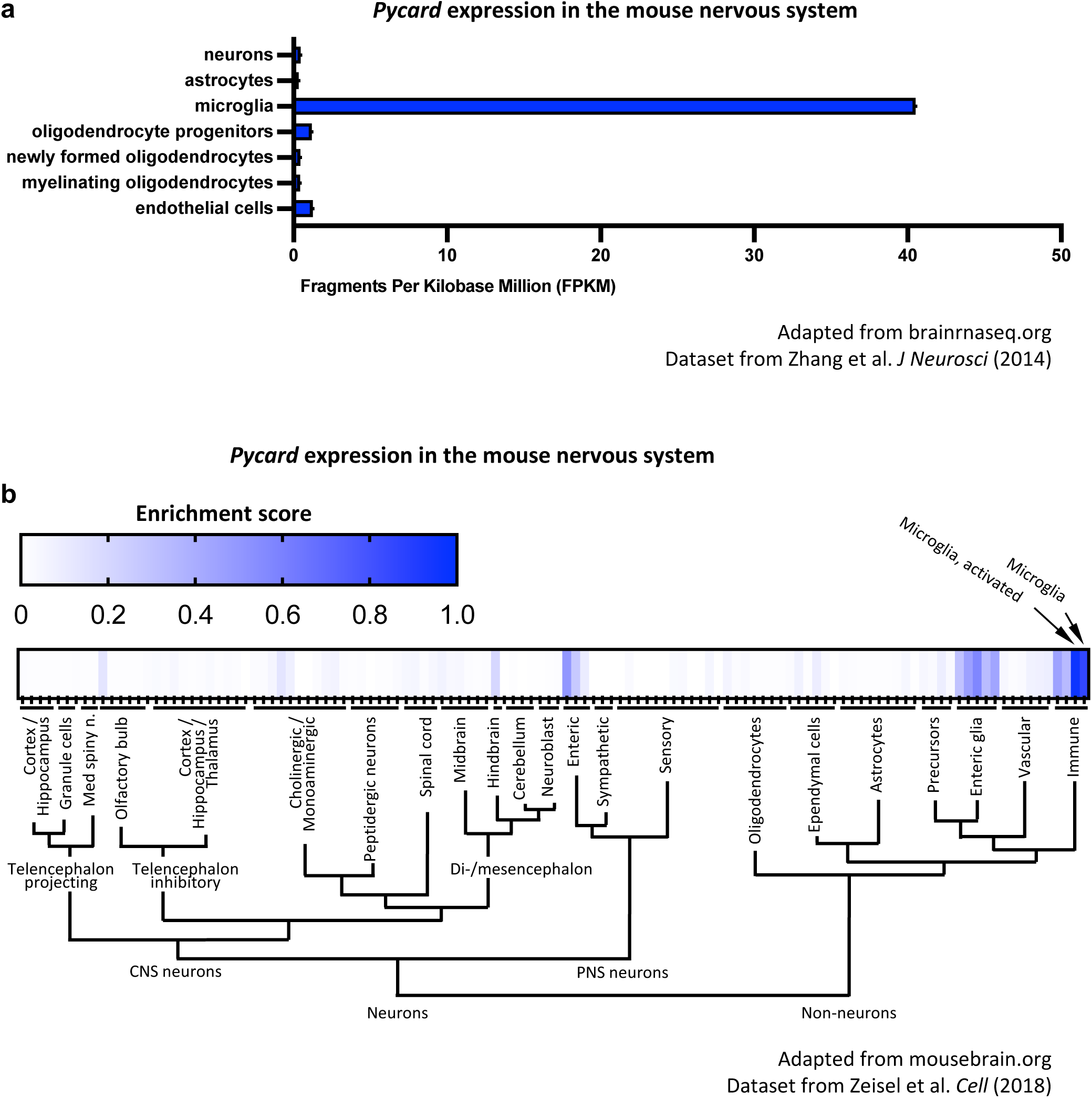
Pycard (alias *Asc*) expression in the mouse nervous system. **(a)** Analysis from brainrnaseq.org, with the dataset from Zhang et al. ^67^ **(b)** Analysis from mousebrain.org, with the dataset from Zeisel et al.^68^

**Suppl. Figure 5.**
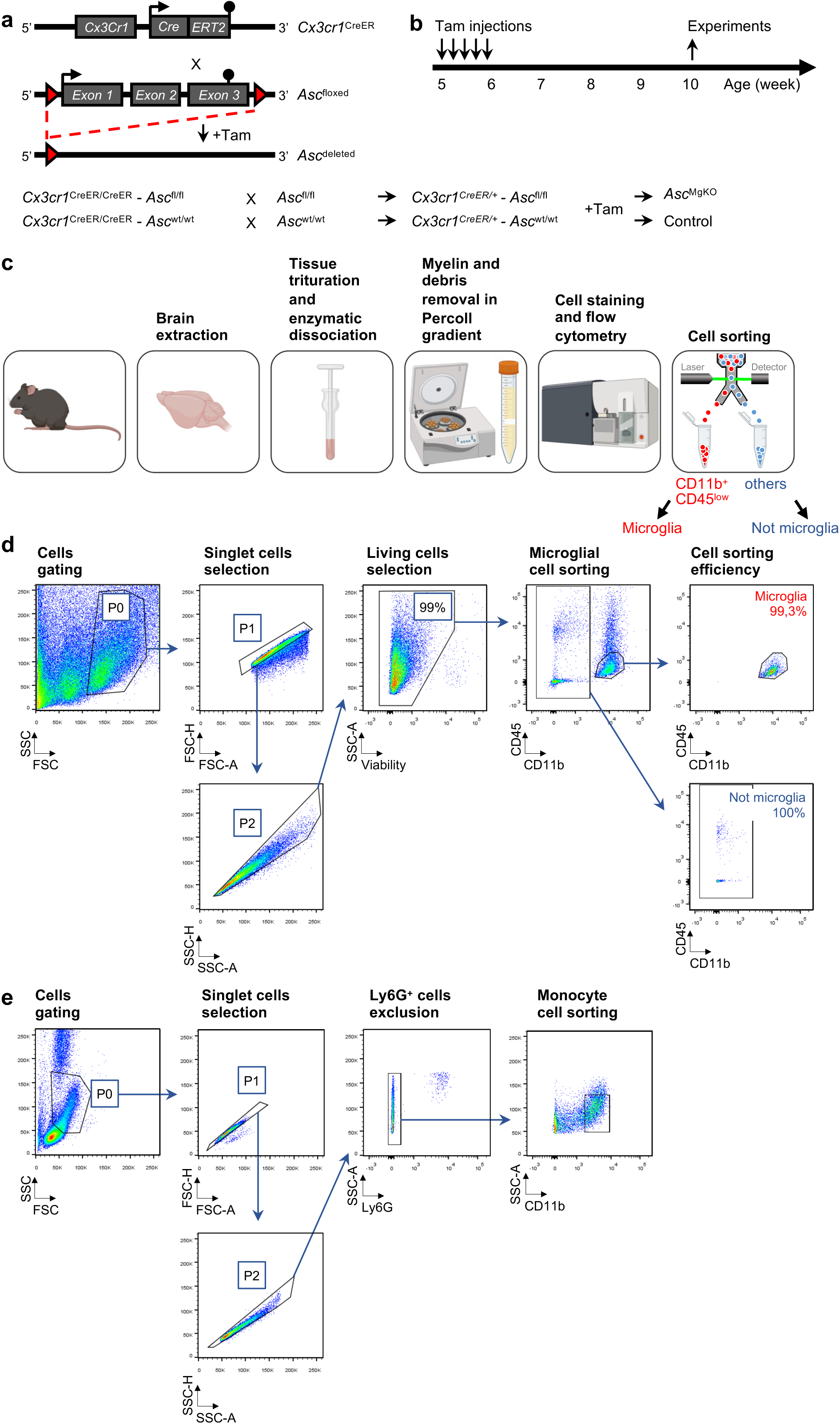
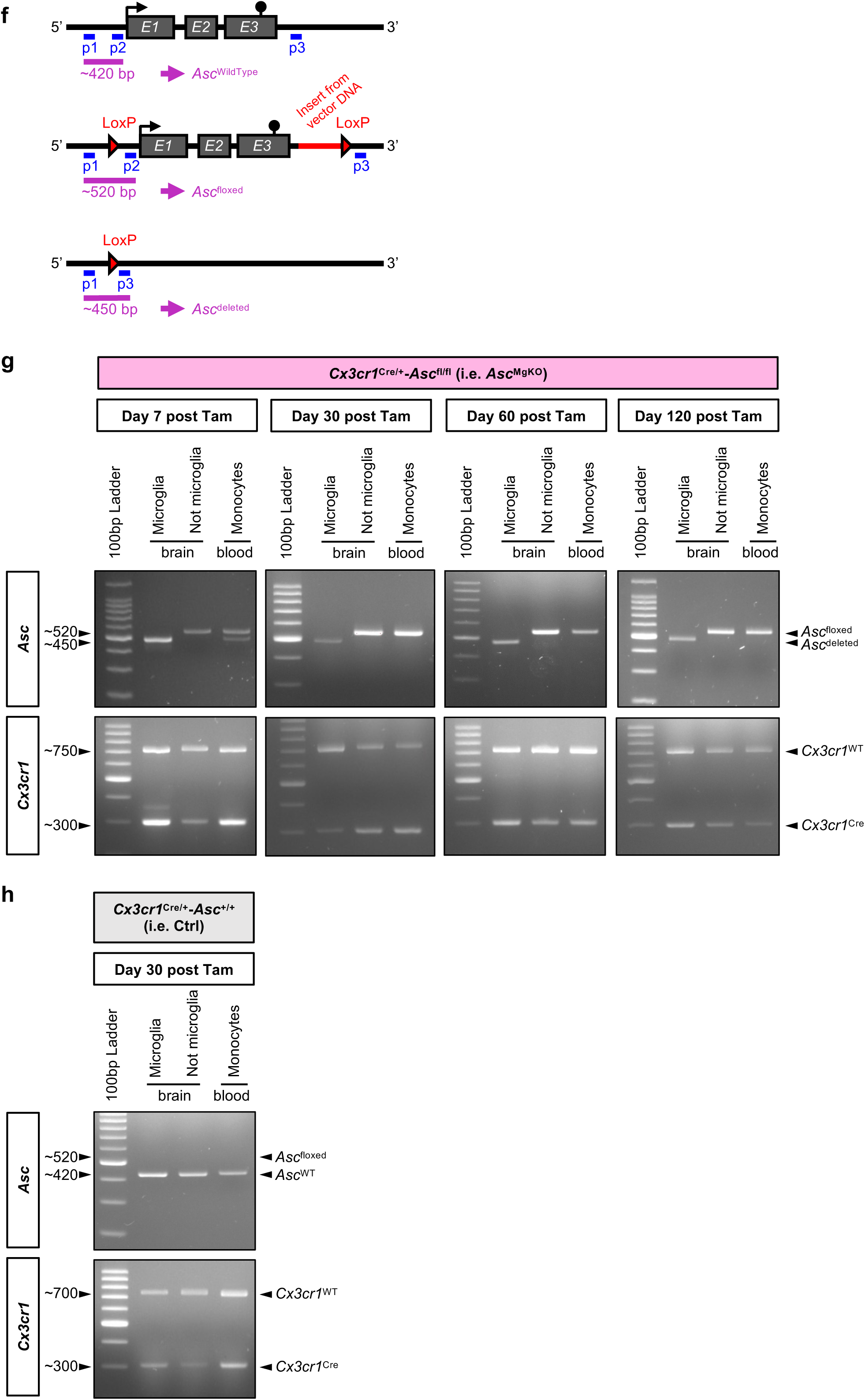
Generation and validation of microglia-specific *Asc* deleted mice (*Asc*^MgKO^). **(a)** Schematic showing the gene targeting strategy used to delete the *Asc* gene in Cx3cr1^+^ cells. Conditional gene deletion was based on the tamoxifen-inducible Cre-lox system driven by the Cx3cr1 promoter. Cx3cr1^CreER^ mice were crossed with mice harboring a floxed *Asc* gene. After tamoxifen treatment, a specific deletion of the *Asc* gene occurred in Cx3cr1^+^ cells. **(b)** Schematic showing protocol of tamoxifen treatment used to induce microglia-specific *Asc* deletion in Cx3cr1^CreER/+^-*Asc*^fl/fl^ mice. 5-week-old mice were injected with tamoxifen (75 mg/kg) for 5 consecutive days. Further experiments were carried out on 10-week-old mice. **(c)** Flowchart of cell purification performed before genomic DNA and total RNA extraction, from cell dissociation to FACS cell sorting. Brain tissues were subjected to enzyme digestion and gentle trituration. Percoll gradient allowed myelin and debris removal. Cell population enriched in microglia was subjected to CD11b and CD45 immunostaining to identify microglia (CD11b^+^/CD45^low^) and not microglial cells (CD11b^-^). **(d)** Gating strategy of FACS-sorted microglia and not microglial cells from brain of C57BL/6J male mice. Cells of interest were localized on side scatter versus forward scatter (SSC and FSC) density plots, corresponding to P0 population. Forward scatter-height (FSC-H) versus forward scatter-area (FSC-A) and side scatter-height (SSC-H) versus side scatter-area (SSC-A) density plots were used to exclude doublets and focus on singlet cells (corresponding to P2 population). Dead cells were excluded by gating on cells negative for a fixable viability stain. Singlet live cells were further sorted using CD11b and CD45 markers. Microglia (CD11b^+^/CD45^low^) were sorted with 99.3% efficiency and not microglial cells (CD11b^-^) were sorted with 100% efficiency. **(e)** Gating strategy of FACS-sorted monocytes from blood of C57BL/6J male mice. After red blood cell removal, remaining blood cells were stained with Ly6G and CD11b markers. Monocytes were gated based on FSC and SSC (P0), followed by singlet cells selection based on FSC-H versus FSC-A (P1) and SSC-H versus SSC-A (P2) density plots. Granulocytes were excluded by gating on cells negative for Ly6G. Monocytes were identified as (Ly6G^-^/CD11b^+^). **(f)** Schematic showing genotyping strategy for *Asc*^WildType^, *Asc*^floxed^, and *Asc*^deleted^ genes (adapted from Drexler et al. ^39^). Exons of the *Asc* targeted locus are shown as dark grey boxes. The targeting vector contains 2 loxP sites to excise the entire *Asc* gene. One forward primer (p1) and two reverse primers (p2 and p3) are shown in blue. The sizes of the amplicons obtained with the triple primer-PCR for *Asc*^WildType^ (∼420 bp), *Asc*^floxed^ (∼520 bp), and *Asc*^deleted^ (∼450 bp) genes are shown in purple. **(g)** Genotyping of microglia-specific *Asc* deleted mice in FACS-sorted microglial cells (CD11b^+^/CD45^low^), not microglial cells (CD11b^-^) from the brain, and FACS-sorted monocytes (Ly6G^-^/CD11b^+^) from the blood of tamoxifen-treated Cx3cr1^Cre/+^-*Asc*^fl/fl^ male mice. Representative agarose gel electrophoresis showing triple primer-PCR analysis performed on DNA extracted from cells at different time points: 7, 30, 60, 120 days after tamoxifen injections. **(h)** Genotyping of control Cx3cr1^Cre/+^-*Asc*^+/+^ mice. Representative agarose gel electrophoresis showing triple primer-PCR analysis performed on DNA extracted from FACS-sorted microglial cells, not microglial cells, and monocytes from Ctrl male mouse 30 days after tamoxifen injections. *Asc*^deleted^ amplicons: band at ∼450 bp. *Asc*^floxed^ amplicons: band at ∼520 bp. Cx3cr1^Cre^: band at ∼300 bp. Cx3cr1^WT^: band at ∼750 bp.

**Suppl. Figure 6.**
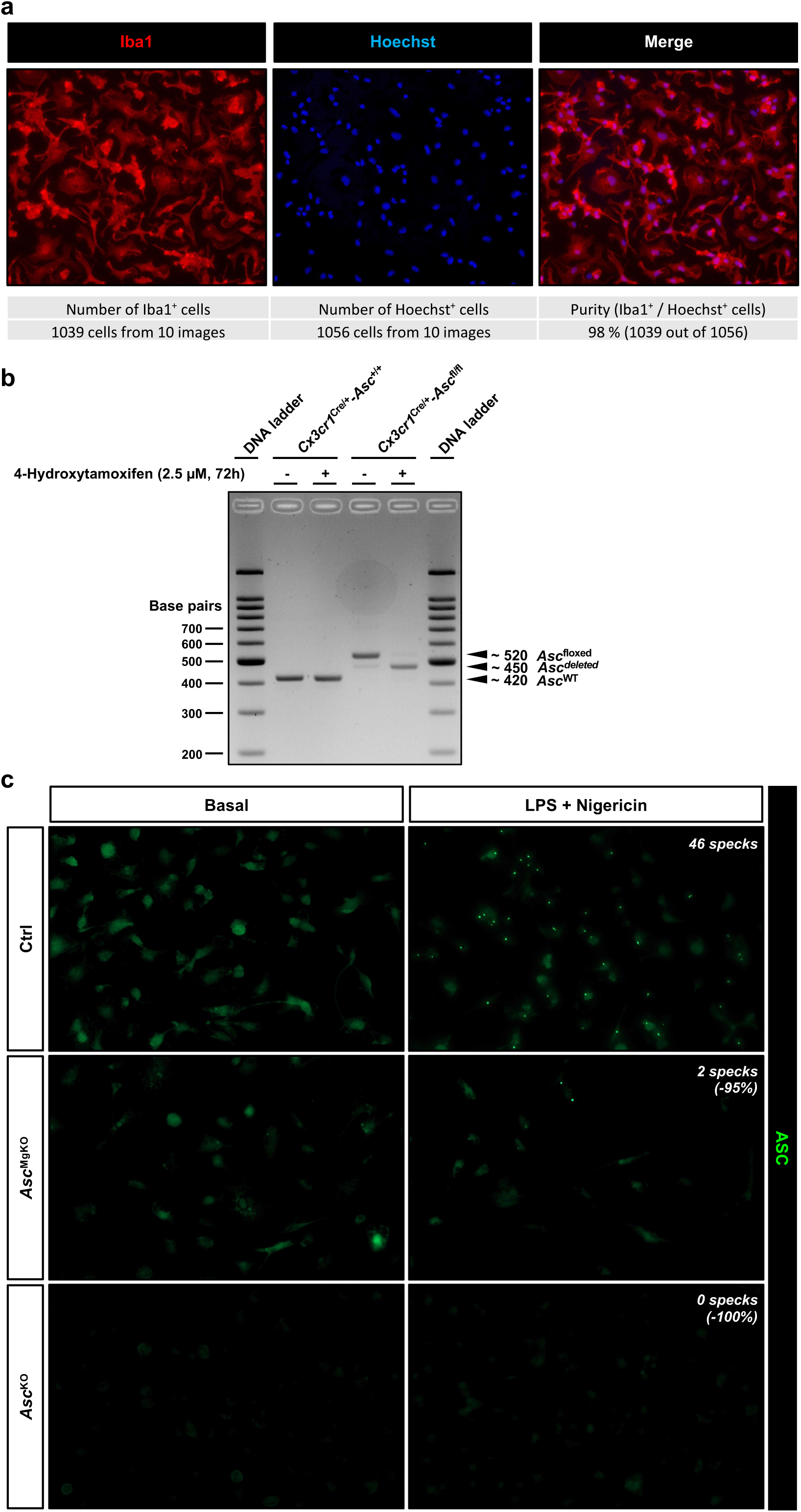
Loss of function in ASC deficient cultured microglia. **(a)** High pure primary culture of Iba1^+^ microglia (>98%) isolated by magnetic cell sorting using CD11b MicroBeads, from dissociated brain cells of newborn C57BL6/J mouse pups. Cells were stained with anti-Iba1 antibody (red) and Hoechst dye (blue). **(b)** Genotyping of primary cultures of Ctrl and *Asc*^MgKO^ microglia. Representative agarose gel electrophoresis showing triple primer-PCR analysis performed on DNA extracted from cells after in vitro tamoxifen treatment (2.5 µM, 72h). **(c)** As shown at high magnification in Figure 4e, this panel shows immunodetected ASC specks in LPS-primed nigericin-treated Ctrl and *Asc*^MgKO^ microglia. Cells were cultured from newborn Cx3cr1^Cre/+^-*Asc*^+/+^ (Ctrl) and Cx3cr1^Cre/+^-*Asc*^lox/lox^ mice and were all treated with 4-hydroxytamoxifen in vitro. After ASC depletion, cells were stimulated with LPS and nigericin, fixed with paraformaldehyde, and stained with anti-ASC antibody (green), anti-iba1 antibody (red), and Hoechst dye (blue). This supplementary panel displays the green channel only and provides a large field at low magnification to reveal the drastic reduction of ASC specks in LPS-nigericin stimulated *Asc*^MgKO^ microglia. Arrowheads show immunolabeled green fluorescent ASC specks. Full *Asc*^KO^ microglia isolated from total ASC^KO^ mice were used as control, showing a complete loss of ASC immunoreactivity.

**Suppl. Figure 7.**
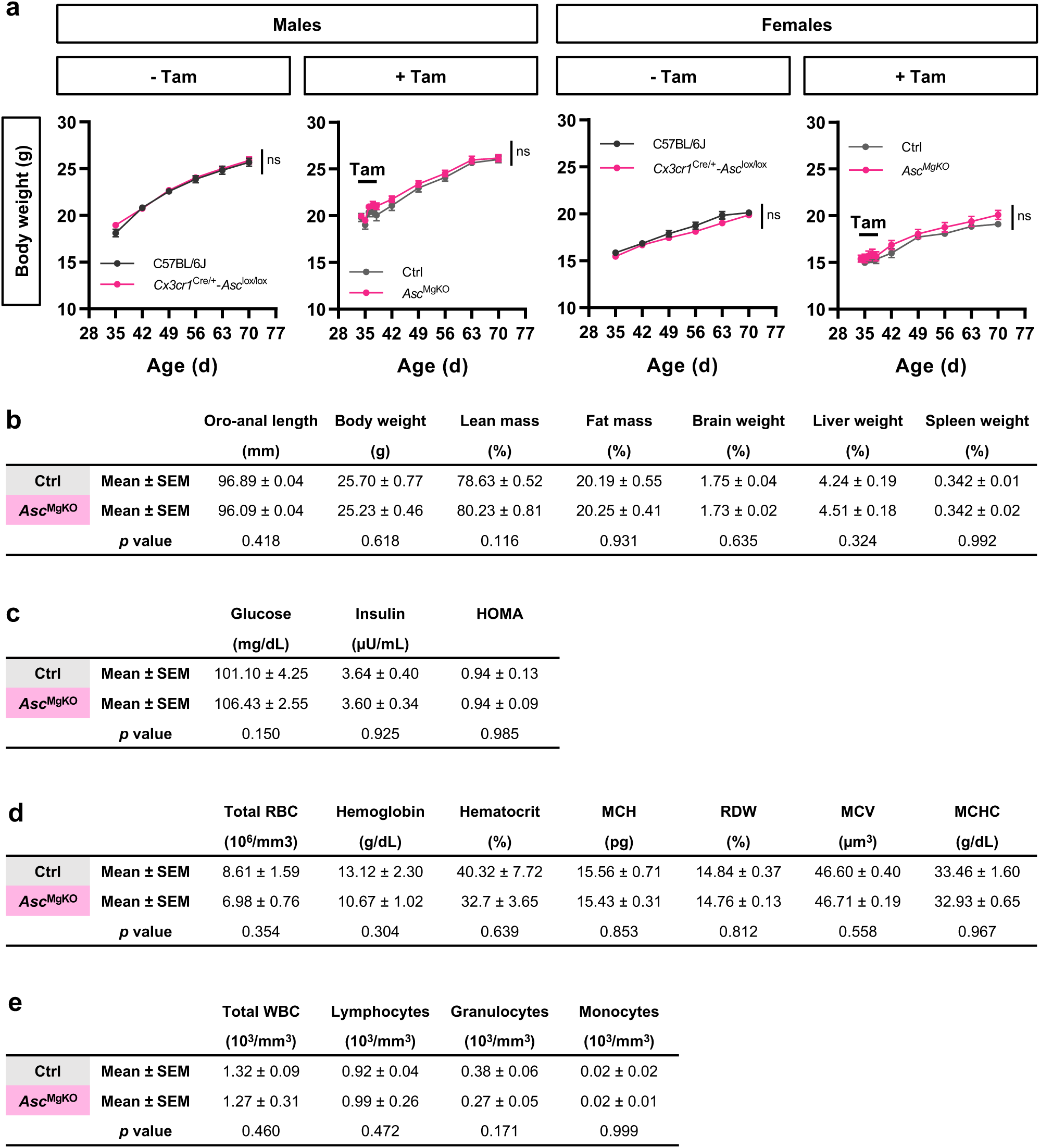
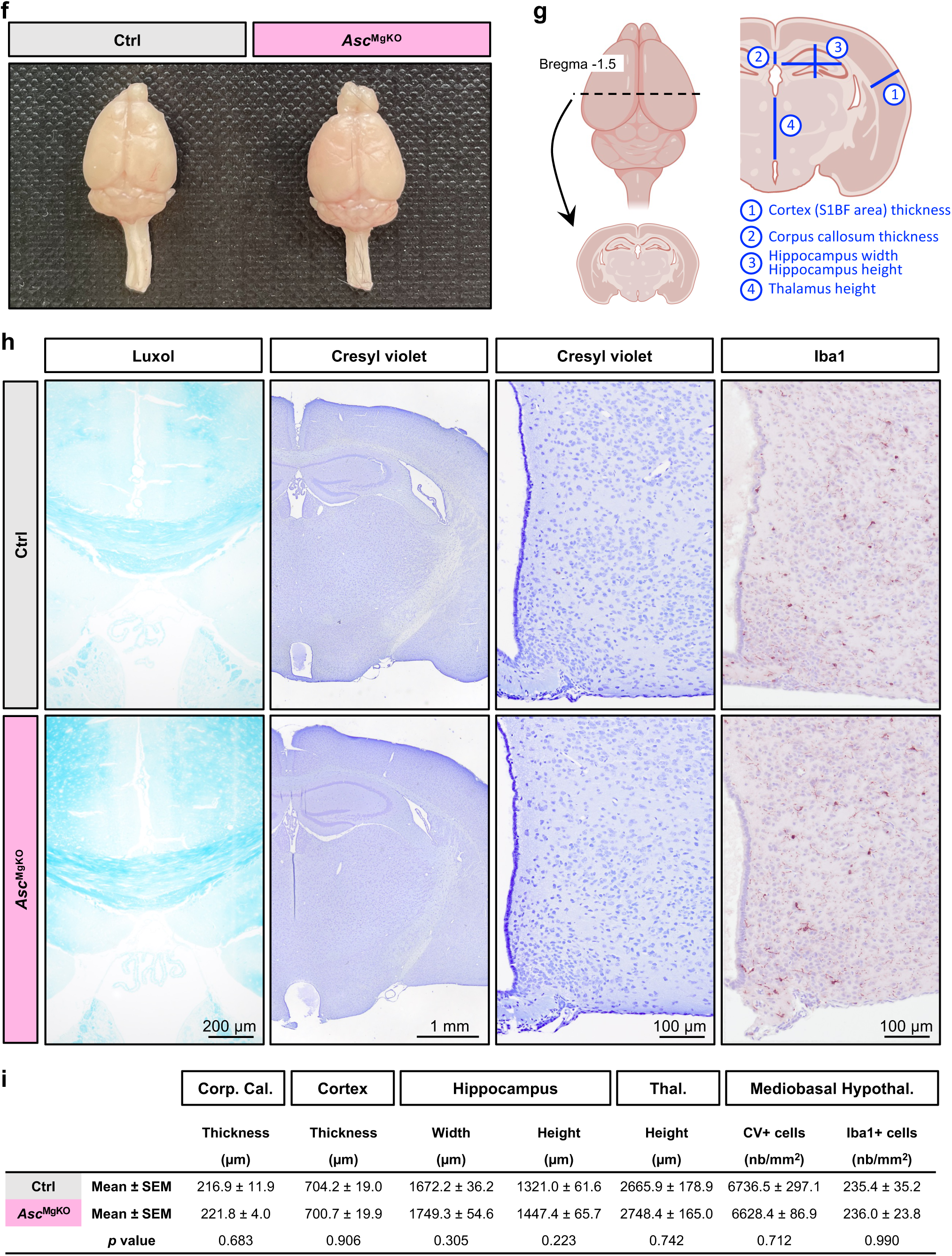
Phenotypic characterization of *Asc*^MgKO^ mice. **(a)** Growth curves of *Asc*^MgKO^ and Ctrl mice, in both sexes, with or without tamoxifen (Tam) treatment, showing no growth deviation on standard diet compared to C57BL/6J mice (males C57BL/6J -Tam: n=16, Cx3cr1^Cre/+^-*Asc*^fl/fl^ -Tam : n=44; males Ctrl: n=24, *Asc*^MgKO^: n=27; females C57BL/6J -Tam: n=15, Cx3cr1^Cre/+^-*Asc*^fl/fl^ -Tam : n=33; females Ctrl: n=8, *Asc*^MgKO^: n=10). **(b)** Size, body weight and composition, and organ weights of 10-week-old male Ctrl and *Asc*^MgKO^ ^mice^, 30 days after tamoxifen treatment, showing no metabolic impairment or malformation on standard diet (Ctrl, n=15; *Asc*^MgKO^, n=15). **(c)** Fasting blood glucose, plasma insulin and HOMA index in 10-week-old male Ctrl and *Asc*^MgKO^ mice, showing no difference in blood glucose control on standard diet (Ctrl, n=15; *Asc*^MgKO^, n=15). (**d,e)** Complete blood count of 10-week-old male Ctrl and *Asc*^MgKO^, 30 days after tamoxifen treatement, showing no difference on standard diet in **(d)** red blood cells, hemoglobin, hematocrit, and others red blood cells parameters including mean corpuscular hemoglobin (MCH), red blood cell distribution width (RDW), mean cell volume (MCV), and mean corpuscular hemoglobin concentration (MCHC); and **(e)** white blood cells (Ctrl, n=5 mice; *Asc*^MgKO^, n=7 mice). **(f)** Picture of two brains dissected from 10-week-old male Ctrl and *Asc*^MgKO^ mice, illustrating the absence of difference in brain size and morphology. **(g)** Schematic for brain sectioning and morphometric parameters used in **(h,i)**. **(h)** Representative images of Luxol staining for corpus callosum thickness determination, Cresyl violet staining for gross morphology and cell density determination, and Iba1 immunostaining for microglia cell counting in brain sections of 11/12-week-old male Ctrl and *Asc*^MgKO^ mice. (i) Statistical analysis revealed no signs of demyelination and malformation, and no change in cell density and microglia density in brain of *Asc*^MgKO^ mice (Ctrl, n= 6 mice; *Asc*^MgKO^, n= 6 mice). **Statistics.** Data are mean ± standard error of the mean (SEM). Asterisks indicate statistical differences between groups: ns, non-significant. Data were analysed using two-way ANOVA (in a), and two-tailed Mann-Whitney test (in b, c, d, e, i).

**Suppl. Figure 8.**
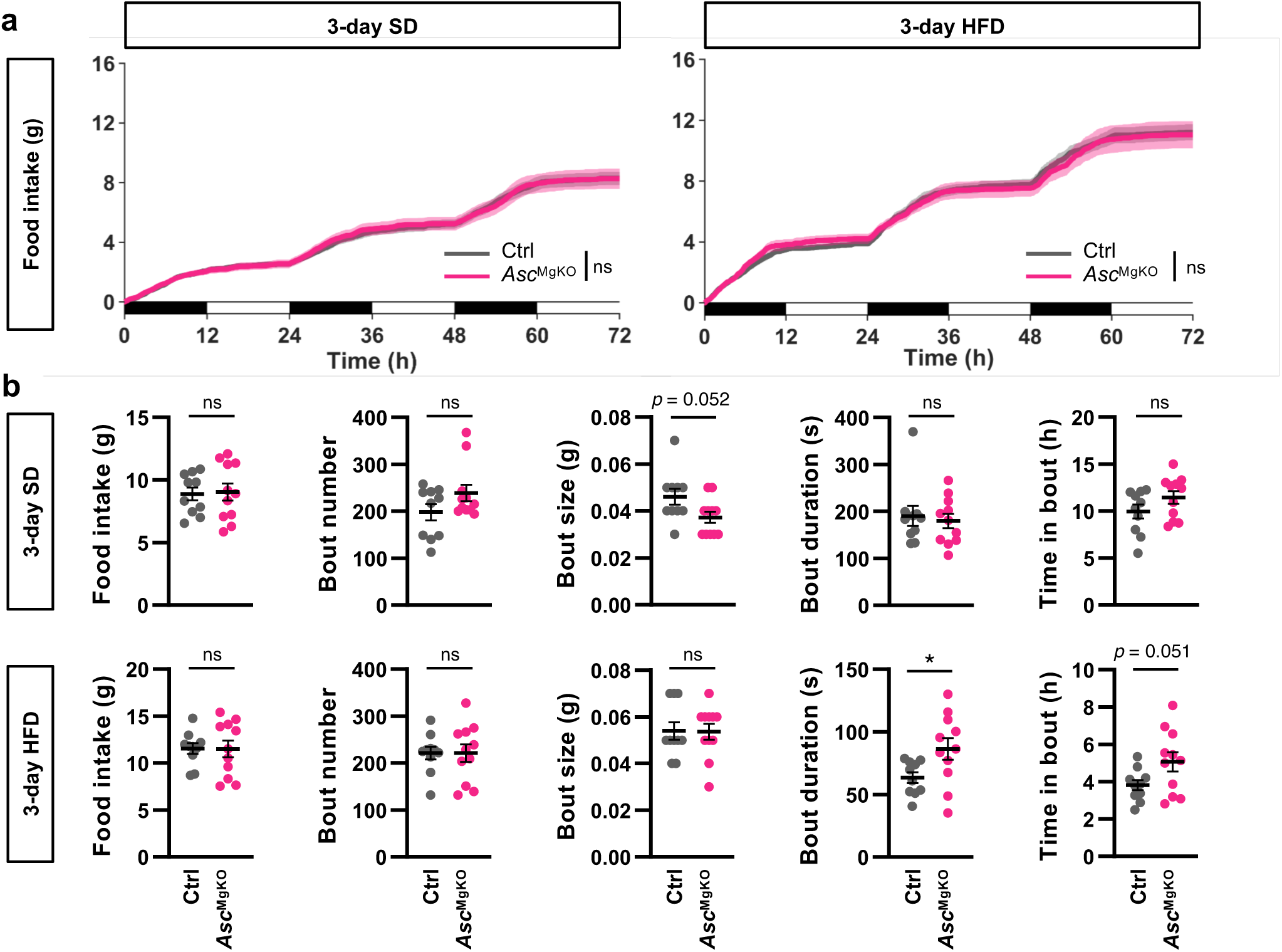
Feeding behaviour in female *Asc*^MgKO^ mice. **(a)** Cumulative food intake of 12/13-week-old female Ctrl and *Asc*^MgKO^ mice fed with SD or HFD ad libitum for 72 hours (Ctrl: n=10 mice; *Asc*^MgKO^: n=11 mice). **(b)** Microstructure of food intake including total food intake, number of feeding bouts, size of feeding bouts, duration of feeding bouts, and percentage of time spent in feeding bouts in female Ctrl and *Asc*^MgKO^ mice fed with SD or HFD ad libitum for 72 hours (Ctrl: n=10 mice; *Asc*^MgKO^: n=11 mice). **Statistics.** Data are mean ± standard error of the mean (SEM). Asterisks indicate statistical differences between groups: *p<0.05; ns, non-significant. Data were analysed using two-way ANOVA and (in a) and two-tailed unpaired Student’s test (in b).

